# Heterozygote Advantage Can Explain the Extraordinary Diversity of Immune Genes

**DOI:** 10.1101/347344

**Authors:** Mattias Siljestam, Claus Rueffler

## Abstract

**Abstract:** The majority of highly polymorphic genes are related to immune functions and with over 100 alleles within a population, genes of the major histocompatibility complex (MHC) are the most polymorphic loci in vertebrates. How such extraordinary polymorphism arose and is maintained is controversial. One possibility is heterozygote advantage (HA), which can in principle maintain any number of alleles, but biologically explicit models based on this mechanism have so far failed to reliably predict the coexistence of significantly more than ten alleles. We here present an eco-evolutionary model showing that evolution can result in the emergence and maintenance of more than 100 alleles under HA if the following two assumptions are fulfilled: first, pathogens are lethal in the absence of an appropriate immune defence; second, the effect of pathogens depends on host condition, with hosts in poorer condition being affected more strongly. Thus, our results show that HA can be a more potent force in explaining the extraordinary polymorphism found at MHC loci than currently recognized.

## Introduction

Heterozygote advantage (HA) is a well-established explanation for single locus polymorphism, with the sickle cell locus as a classical text book example (***Allison, 1954***). However, whether HA is generally important for the maintenance of genetic polymorphism is questioned (***Hedrick, 2012***; ***Sellis et al., 2016***). Genes of the major histocompatibility complex (MHC), responsible for inducing immune defences by recognizing the agretopes of the pathogenic antigens, are the most polymorphic loci among vertebrates (***Duncan et al., 1979***; ***Apanius et al., 1997***; ***Penn, 2002***; ***Sommer, 2005***; ***Eizaguirre and Lenz, 2010***). HA as an explanation for this high level of polymorphism was introduced almost 50 years ago by ***Doherty and Zinkernagel*** (***1975***). The idea suggests that individuals with MHC molecules from two different alleles are capable of recognizing a broader spectrum of pathogens, resulting in higher fitness. This is especially evident when the MHC molecules of the two alleles have complementary immune profiles (***Pierini and Lenz, 2018***), a phenomenon known as divergent allele advantage (***Wakeland et al., 1990***) and ***Stefan et al. (2019***) show that this allows for the coexistence of alleles with larger variation in their immune efficiencies. Early theoretical work suggested that HA can maintain an arbitrarily high number of alleles if these alleles have appropriately fine-tuned homo- and heterozygote fitness values (***Kimura and Crow, 1964***; ***Wright, 1966***; ***Maruyama and Nei, 1981***). However, later work suggests that such genotypic fitness values are unlikely to emerge through random mutations (***Lewontin et al., 1978***). More mechanistic models have also failed to reliably predict very high allele numbers (***Spencer and Marks, 1988***; ***Hedrick, 2002***; ***de Boer et al., 2004***; ***Borghans et al., 2004***; ***Stoffels and Spencer, 2008***; ***Trotter and Spencer, 2008, 2013***; ***Ejsmond and Radwan, 2015***; ***Lau et al., 2015***). As a result, HA plays only a minor role in current explanations of polymorphism at MHC loci (***Hedrick, 1999***; ***Gould et al., 2004***; ***Wegner, 2008***; ***Kekäläinen et al., 2009***; ***Eizaguirre and Lenz, 2010***; ***Lenz, 2011***; ***Loiseau et al., 2011***), despite empirical evidence for its existence (***Doherty and Zinkernagel, 1975***; ***Hughes and Nei, 1989***; ***Jeffery and Bangham, 2000***; ***Penn et al., 2002***; ***McClelland et al., 2003***; ***Froeschke and Sommer, 2005***; ***Kekäläinen et al., 2009***; ***Oliver et al., 2009***; ***Lenz, 2011***). Consequently, other mechanisms are suggested to be important for the maintenance of allelic diversity, such as Red-Queen dynamics, fluctuating selection and disassortative mating (***Apanius et al., 1997***; ***Hedrick, 1999***; ***Penn, 2002***; ***Borghans et al., 2004***; ***Wegner, 2008***; ***Spurgin and Richardson, 2010***; ***Loiseau et al., 2011***; ***Ejsmond and Radwan, 2015***; ***Ejsmond et al., 2023***).

Our study challenges this status quo by demonstrating that HA is a potent force that can drive the evolution and subsequent maintenance of more than 100 alleles. To demonstrate that it is indeed heterozygote advantage that is responsible for allelic diversity in our model, we deliberately keep all aspects of the pathogen community fixed to exclude any Red-Queen dynamics. The novelty of our approach lies in the fact that we do not rely on hand-picked genotypic fitness values. Instead, these fitness values emerge from our eco-evolutionary models, where the allelic values that allow for extraordinary polymorphism are found by evolution in a self-organized process. We do not claim that HA is the only mechanism responsible for the diversity of MHC alleles in nature. However, our results show that HA can be more important than currently recognized.

## Model

We investigate the evolution at an MHC locus using mathematical modelling and computer simulations. In the following sections, we describe how genotypes map to immune response and ultimately to survival, followed by a description of our evolutionary algorithm.

We assume that the MHC molecules produced by the two alleles at a diploid MHC locus determine the immune response based on antigen recognition against multiple pathogens present in the environment. Our approach is based on the following two key assumptions regarding the relationship between pathogen virulence and host fitness:

a. Virulent pathogens are lethal in the absence of an appropriate immune defence.
b. The effect of pathogens on host survival depends on host condition, with hosts in poorer condition being affected more strongly.

An implication of the second assumption is that the combined effect of multiple pathogens on host survival exceeds the sum of the effects of each pathogen alone.

To incorporate these two assumptions, we assume that the effect of pathogen attacks on host survival acts through the intermediary step of the host’s ‘condition’, which is a proxy for a suit of measurements describing an individual’s body composition and physiology (***Wilder et al., 2016***). In the absence of an adequate immune response, a pathogen attack reduces the condition of a host to zero, causing its death (assumption a). More generally, we assume that the probability to survive is an increasing function of condition and that a host clearing a pathogen is in a weaker condition afterwards. Since the survival probability cannot exceed one, the function that maps condition to survival has to be saturating. Consequently, for high values of conditions, where the survival function has saturated, pathogens reducing condition have small effects on survival. As condition decreases, pathogen induced reductions have larger effects on survival (assumption b). A natural biological intuition for assumption (b) can be drawn from examples like COVID-19 or influenza, where it is well known that these pathogens do not pose a high mortality risk to individuals in good condition, but can significantly increase mortality risk for individuals in poor condition (***Thompson et al., 2004***; ***Zhou et al., 2020***).

A further assumption of our model is the existence of a trade-off between the efficiencies of MHC molecules to induce a defence against different pathogens. Thus, no MHC molecule can perform optimally with respect to all pathogens and an improved efficiency against one set of pathogens can only be achieved at the expense of a decreased efficiency against another set of pathogens. Under such trade-offs, an MHC molecule can be specialized to detect a few pathogens with high efficiency, or, alternatively, be a generalist molecule that can detect many pathogens but with low efficiency. There is empirical support for the existence of such trade-offs. First, many MHC molecules can detect only a certain set of antigens (***Wakeland et al., 1990***; ***Froeschke and Sommer, 2012***; ***Eizaguirre et al., 2012***; ***Chappell et al., 2015***; ***Pierini and Lenz, 2018***) and therefore provide different degrees of protection against different pathogens (***Wakeland et al., 1990***; ***Apanius et al., 1997***; ***Eizaguirre and Lenz, 2010***; ***Froeschke and Sommer, 2012***; ***Eizaguirre et al., 2012***; ***Cortazar-Chinarro et al., 2022***). Second, it has also been found that specialist MHC molecules are expressed at higher levels at the cell surface while generalist MHC molecules that bind less selectively are expressed at lower levels (***Chappell et al., 2015***), potentially to reduce the harm of binding self-peptides. This could explain the lower efficiency of generalist MHC molecules.

We employ two approaches to model this trade-off. First, we use unimodal functions to model the match between MHC molecules and pathogens. This approach has a long history in evolutionary ecology (e.g. ***Levins, 1968***; ***Sheftel et al., 2018***), and, when using Gaussian functions, the model becomes amenable to mathematical analysis. We envisage that these pathogen optima represent distinct pathogen species from diverse taxonomic groups such as fungi, viruses, bacteria, protists, helminths, and prions, among others (***Schmid-Hempel, 2021***). Hence, we expect these pathogen optima to remain approximately constant over the time scales considered in our model. By keeping all aspects of the pathogen community fixed, we exclude Red Queen dynamics and ensure that the observed allelic polymorphism is driven solely by HA.

To demonstrate that the allelic diversity evolving in the Gaussian model does not dependent on the specificities of this model but rather results from the model fulfilling the above assumptions (a) and (b), we implement an alternative and more mechanistic approach to model pathogen recognition. Inspired by ***Borghans et al. (2004***), in this approach, while keeping assumptions (a) and (b) intact, immune defence is based on the match between two binary strings (or bit-strings), one representing the MHC molecule and the other a peptide of the pathogen. In this model, a single MHC allele has the potential to detect several pathogens, which could be interpreted as the different pathogens being more closely related.

By explicitly modelling MHC efficiencies against various pathogens—rather than assuming a fixed proportion of pathogens detected per MHC molecule (as, e.g., ***de Boer et al., 2004***)— our model accounts for the possibility that MHC molecules can have complementary immune profiles. When paired, complementary alleles produce fit heterozygotes able to detect an increased number of pathogen peptides (***Pierini and Lenz, 2018***), exemplifying the concept of divergent allele advantage in the sense of ***Wakeland et al. (1990***).

### Gaussian Model

In this approach, we use Gaussian functions to model the ability of MHC molecules to recognize *m* different pathogens, as illustrated in ***Figure 1***. Here, MHC-alleles and pathogens are represented by vectors ***x*** = (*x*_1_, *x*_2_, …, *x*_*h*_) and ***p***_*k*_ = (*p*_1*k*_, *p*_2*k*_, …, *p*_*hk*_), respectively. The MHC-alleles code for MHC-molecules, and the ability of an MHC molecule to recognize the *k*th pathogen is maximal if ***x*** = ***p***_*k*_. This ability decreases with increasing distance between ***x*** and ***p***_*k*_. The decrease is modelled using an *h*-dimensional Gaussian function *e*_*k*_(***x***), as detailed in ***Equation S14***. The nature of the trade-off can be varied by adjusting the positions of the pathogen optima and the shape of the Gaussian functions.

**Figure 1.**
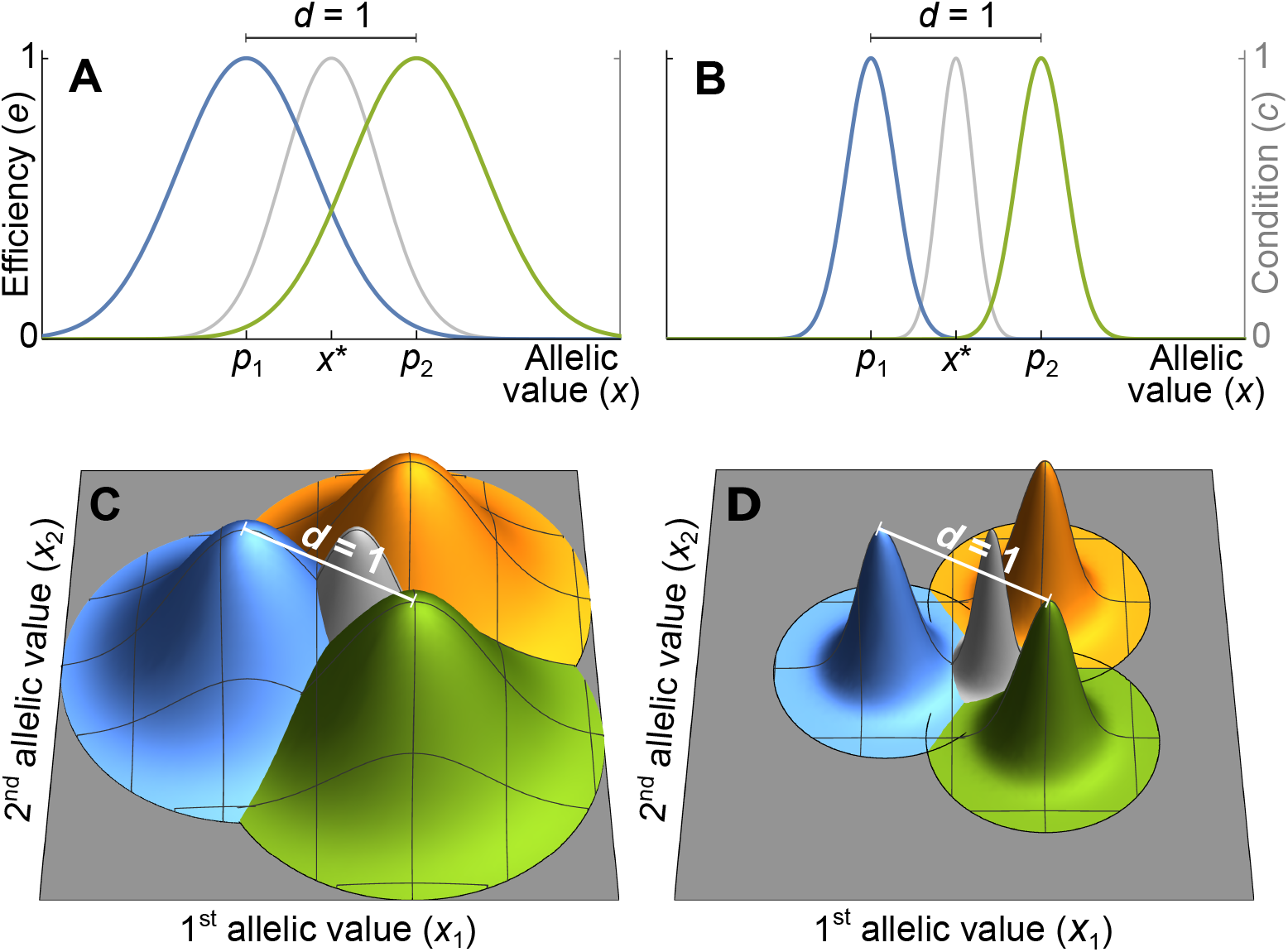
Efficiency against two pathogens (coloured lines in ***A-B***) and three pathogens (coloured cones in ***C-D***) as a function of allelic values ***x***. Efficiencies are modelled with Gaussian functions with pathogen optima at equal distances *d* = 1 (indicated by *p*_1_ and *p*_2_ in ***A, B***). The width of the Gaussian functions, which determine how severely pathogens affect hosts with suboptimal MHC molecules, is given by the virulence parameter *v*. With high virulence (*v* = 7, narrow Gaussians in ***B, D***), alleles away from the optima have a low efficiency, while for a low virulence (*v* = 2.5, wide Gaussians in ***A, C***) efficiency is higher. Gray lines and cones give the condition *c* of homozygote individuals. The generalist allele, maximizing condition, is located at the centre with equal distance to all pathogen optima (indicated by *x*^*^ in ***A, B***).

Without loss of generality, we can reduce the dimension of the vectors ***x*** and ***p***_*k*_ to *h* = *m* − 1, such that ***x*** = (*x*_1_, *x*_2_, …, *x*_*m*−1_) and ***p***_*k*_ = (*p*_1*k*_, *p*_2*k*_, …, *p*_*m*−1,*k*_). For example, in ***Figure 1A, B***, where *m* = 2, the x-axis represents the unique line passing through two pathogen optima in a trait space of potentially much higher dimension. Similarly, in ***Figure 1C, D***, where *m* = 3, the two-dimensional coordinate system represented by the gray surfaces describes the unique plane passing through three pathogen optima. Mathematically speaking, *m* linearly independent pathogen optima form the basis of a vector space of dimension *m* − 1, which we choose as the coordinate system for the vectors ***x*** and ***p***. Allelic vectors outside this set are necessarily maladapted for all pathogens along at least one dimension, and owing to our dimensionality reduction we ignore such trait vectors.

We examine two versions of the Gaussian model. The first one is based on two symmetry assumptions and shown in ***Figure 1***: pathogen optima are placed symmetrically such that the distance between any two pathogens equals 1, and the Gaussian functions *e*_*k*_(***x***) are isotropic (rotationally symmetric) and of equal width. This allows to simplify the covariance matrix in the Gaussian function *e*_*k*_(***x***) (***Equation S14***) such that it can be replaced with a single parameter *v* (***SI Appendix 6.2***),

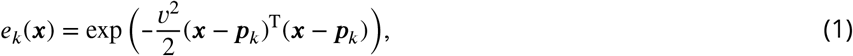

where the superscript T indicates vector transposition. The parameter *v*, to which we refer as virulence, is the inverse of the width of the Gaussian function. If the Gaussian function is narrow, corresponding to a high virulence *v*, a pathogen causes significant harm if MHC-molecules are not well adapted against it (***Figure 1B*** and ***D***). On the other hand, if the Gaussian function is wide, corresponding to a low virulence *v*, a pathogen causes less harm (***Figure 1A*** and ***C***).

We relax these symmetry assumptions in the second version, where we allow for Gaussian functions with arbitrary shape and position. Since the results for the two versions are similar, we here focus on the case with symmetry and refer to ***SI Appendix 4, 6.2*** and ***7*** for results based on general Gaussian functions.

### Bit-String Model

Our second approach is inspired by ***Borghans et al. (2004***) and commonly referred to as a bit-string model. Pathogens are assumed to produce *n*_pep_ peptides, and for a pathogen to cause virulence, all of its peptides have to avoid detection by the host’s MHC molecules. We here equate MHC-alleles with the MHC-molecule they code for, and both MHC molecules and pathogen peptides are represented by binary strings (or bit-strings) of, following ***Borghans et al. (2004***), length 16.

The probability that an MHC molecule detects a pathogen peptide increases with the maximum match length of consecutive matches between their binary strings. For an MHC molecule ***x*** and the *k*th peptide of the *i*th pathogen, this match length is denoted *L*_*ki*_(***x***), or *L* for short (see ***Figure 2A***). The corresponding detection probability, denoted *D*(*L*_*ki*_(***x***)), is then given by the logistic function

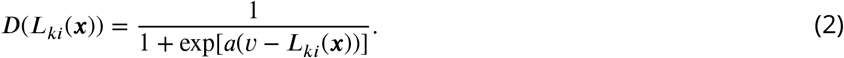

**Figure 2.**
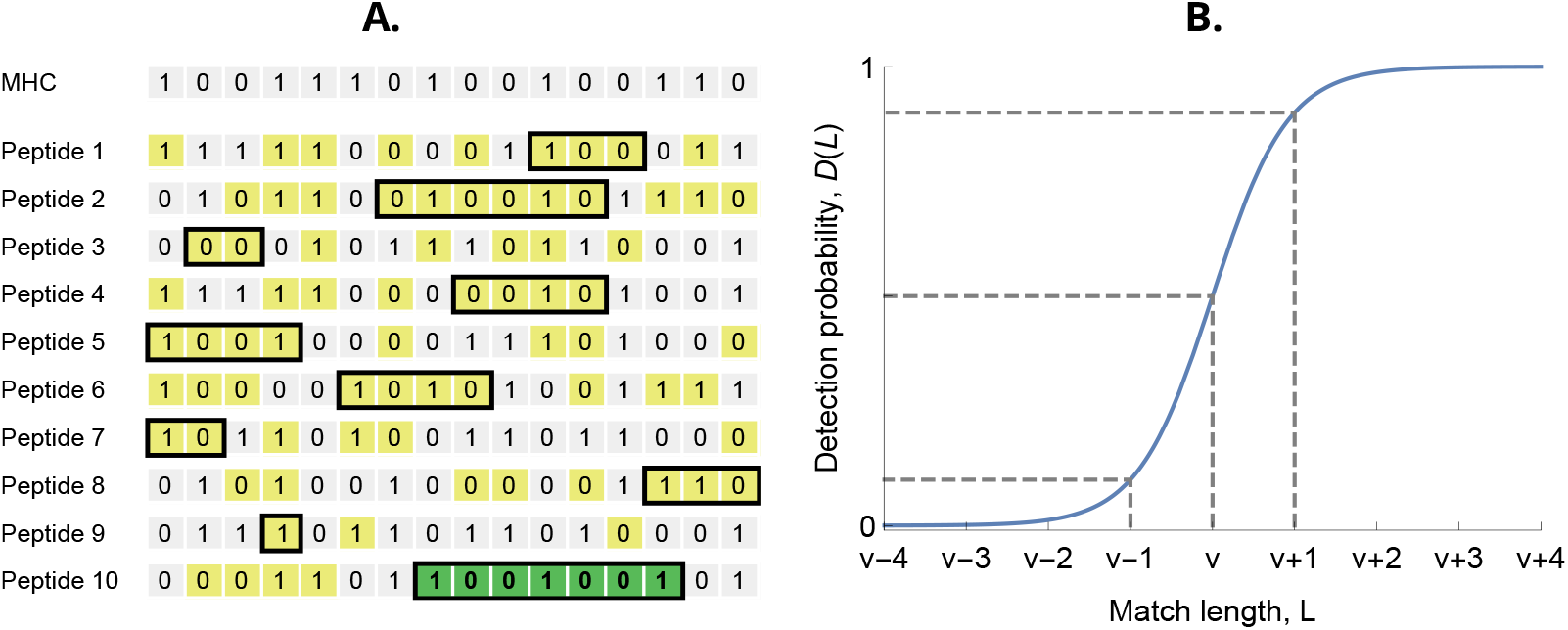
(**A**) MHC bit-string matching against a pathogen with *n*_pep_ = 10 peptides. Yellow indicates a match between MHC and peptide bits. The longest consecutive match per peptide (*L*) is indicated with a black box. The longest match over all peptides occurs for the last peptide, marked in green, with match length *L* = 7. (**B**) Detection probability for peptides as a function of match length *L* (***Equation 2*** with *a* = log(9)). The dashed lines indicate, from left to right, 10%, 50% and 90% detection probability.

Here, *v* denotes the required match length *L* for a 50% chance of detection. The parameter *v* has again the interpretation of virulence, with higher values indicating pathogen peptides that are harder to detect by MHC molecules. The positive parameter *a* governs the steepness of the function *D*. We choose *a* = log(9), which results in *D*(*L*) equalling 10% when *L* = *v* − 1 and 90% when *L* = *v* + 1 (***Figure 2B***). Finally, the realised efficiency of an MHC molecule ***x*** against the *k*th pathogen is given by the probability of detecting at least one of its *n*_pep_ peptides, which equals

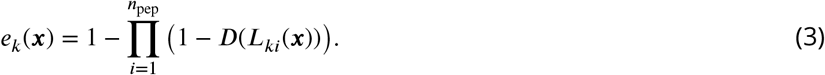

### From Immune Defence to Survival

For both versions of our model, we assume that MHC alleles are co-dominantly expressed (***Eizaguirre and Lenz, 2010***; ***Abbas et al., 2014***), and an individual’s efficiency to recognize pathogens of type *k* is given by the arithmetic mean of the efficiencies from its two alleles. We want to note that assuming co-dominance gives more conservative results in terms of the number of coexisting alleles, as dominance would increase the degree of heterozygote advantage.

For each pathogen attack, an individual’s condition *c* is reduced by a certain fraction that depends on the efficiency of the defence *e* against that pathogen. Since each individual is exposed to all pathogens during their lifetime, the condition *c* is determined by the product of its defences against all pathogens,

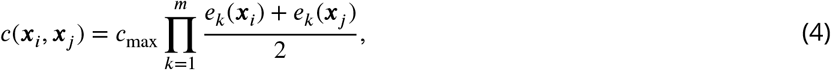

where ***x***_*i*_ and ***x***_*j*_ represent the MHC alleles the host carries at the focal locus, and *c*_max_ is the condition of a hypothetical individual with perfect defence against all pathogens (see ***Supplementary 6.2*** for more details). Because *e*_*k*_(***x***) *<* 1, condition is reduced with each additional pathogen in a proportional manner. The multiplicative nature of ***Equation 4*** has the effect that a poor defence against a single pathogen is sufficient to severely compromise condition, and therefore survival (see next paragraph), fulfilling assumption (a) above.

Finally, survival *s* is an increasing but saturating function of an individual’s condition *c*,

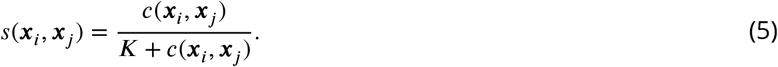

Here, ***K*** is the survival half-saturation constant, giving the condition *c* required for a 50% chance of survival. This function fulfils assumption (b) above as long as ***K*** is not too large. Individuals in good health then have a condition *c* far above ***K***, and a decrease in condition only has a small effect on survival. If *c* is lower than ***K***, then the host is in bad health and any additional pathogen causes a large reduction in survival *s* (orange lines in bottom panel of ***Figures 3*** and ***6***).

**Figure 3.**
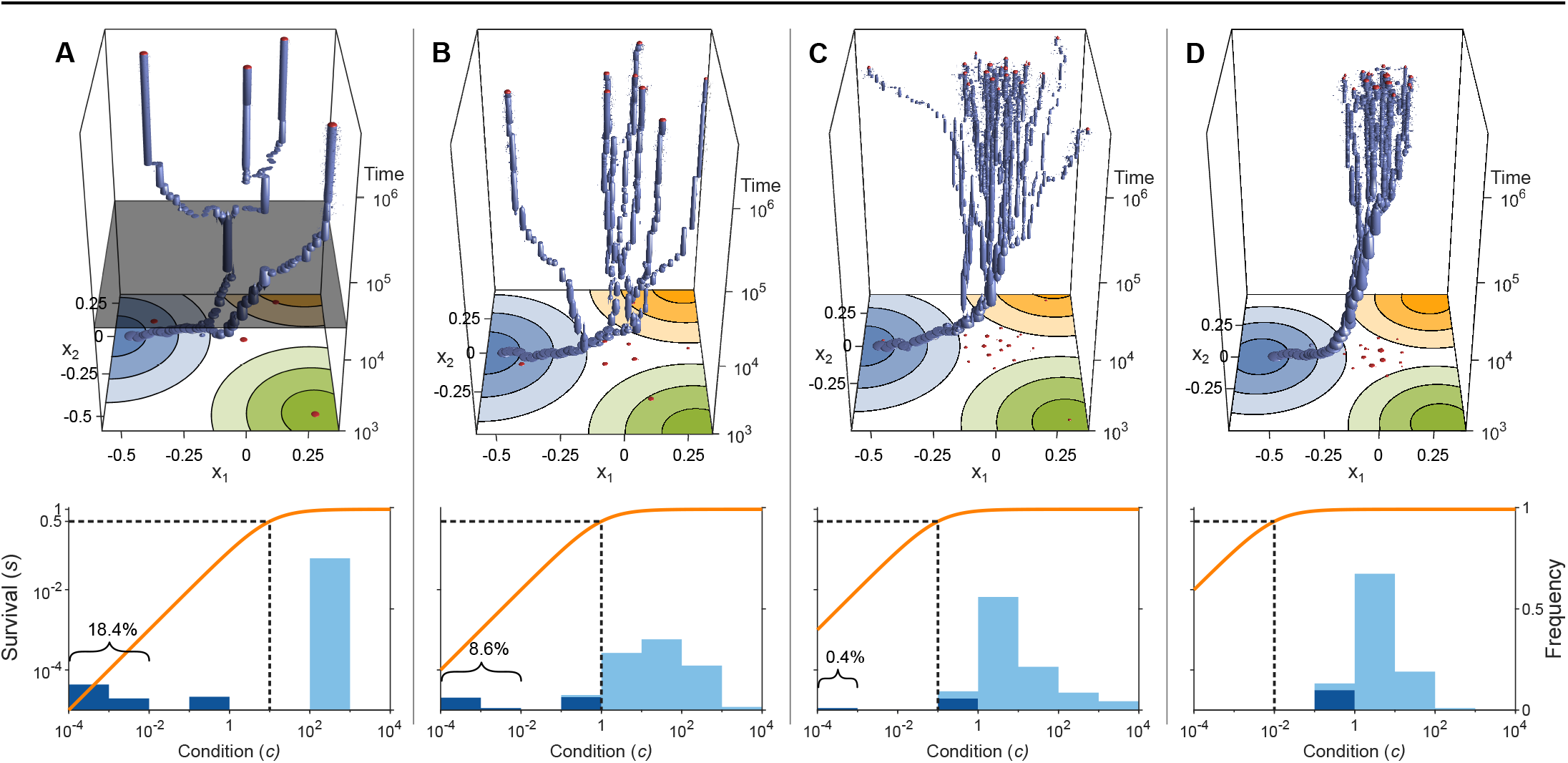
Evolution of allelic values under the Gaussian model in the presence of three pathogens (arranged as in ***Figure 1D***) for four different values of the survival half-saturation constant *K* (***A***: *K* = 10, ***B***: *K* = 1, ***C***: *K* = 0.1, ***D***: *K* = 0.01; dashed line in lower panel). The top panel shows individual-based simulations. The two horizontal axes give the two allelic values ***x*** = (*x*_1_, *x*_2_) that characterize an allele, while the vertical axis shows evolutionary time. The thickness of the blue tubes is proportional to allele frequencies. Allelic values at the last generation are projected as red dots on the top as well as on the bottom plane. Coloured circles represent the contour lines of the Gaussian efficiency functions *e*_*k*_(***x***) shown in ***Figure 1D***. In all simulations, gradual evolution leads toward the generalist allele ***x***^*^ = (0, 0) and branching occurs in its neighbourhood, as predicted by our analytical derivations (***SI Appendix 7.1***). In ***A*** there are three consecutive branching events with the second branching event marked by the grey plane (*n*_e_ = 4.0; for details regarding *n*_*e*_, see the legend of ***Figure 4***). ***B*** and ***C*** show that, as *K* decreases, the number of branching events increases, resulting in more coexisting alleles (*n*_e_ = 7.8 and *n*_e_ = 16.5, respectively). Finally, ***D*** reveals that, as *K* decreases even further such that already low condition values result in high survival, the number of branching events decreases again, resulting in a set of alleles closely clustered around the generalist allele (*n*_e_ = 10.2). The bottom panel shows survival *s* as a function of condition *c* as defined by ***Equation 5*** on a log-log scale (orange line, left vertical axis) and the frequencies of individual conditions at the final generation (dark blue bars for homozygotes and light-blue bars for heterozygotes, right vertical axis; conditions from 0 to 10^−4^ are incorporated into the first bar). These panels show that increased allelic diversity results in a lower proportion of homozygote individuals, which have lower survival. Other parameter values: *v* = 7, *N* = 2 × 10^5^, *µ* = 10^−6^, and *δ* = 0.016.

In summary, ***Equation 4*** and ***5*** entail that assumptions (a) and (b), as formulated above, are satisfied. Using two distinct models to describe the interaction of hosts and pathogens, which both impose a trade-off between the ability to detect different pathogens—namely the Gaussian and the bit-string model—we demonstrate below that HA emerges as a potent force capable of driving the evolution of a very high number of coexisting alleles.

## Analysis

To study the evolutionary dynamics of allelic values ***x*** in both the Gaussian and the bit-string model, we simulate a diploid Wright-Fisher model with mutation and selection (***Fisher, 1930***; ***Wright, 1931***). Thus, we consider a diploid population of fixed size *N* with non-overlapping generations and random mating. Individuals produce, independent of their genotype, a large number of offspring, resulting in deterministic Hardy-Weinberg proportions before viability selection. After viability selection, which is based on ***Equation 5*** and adjusts the proportion of genotypes accordingly, stochasticity is introduced by random multinomial sampling of *N* surviving offspring, which constitute the adult population of the next generation. Using this model, we follow the fate of recurrent mutations that occur with a per-capita mutation probability *µ*. The long-term evolutionary dynamics is obtained by iterating this procedure (***Figure 3***, top panel) until the number of alleles equilibrates. This procedure can result in high numbers of coexisting alleles, where the emerging allelic polymorphism is driven by increasing the alleles’ expected survival (or marginal fitness, see ***Equation S5-S6*** in ***SI Appendix 6.1***).

For the Gaussian model, mutations are drawn from an isotropic normal distribution with expected mutational effect-size *δ* (***SI Appendix 3***). We here focus on mutations of small effect (*δ* = 0.016 in ***Figure 3*** and *δ* = 0.03 in ***Figure 4***) and thus near-gradual evolution. The effect of smaller and larger effect sizes is investigated in ***SI Appendix 3***. To minimize computation time, simulations (other than those in ***Figure 3***) are initialized at the trait vector that is given by the mean of the vectors describing the pathogens. In the bit-string model, the *m* pathogens are each given *n*_pep_ randomly drawn bit-strings at the beginning of a simulation and the host population is initialized with a single MHC allele given by a randomly drawn bit-string. Mutations change a random bit of the MHC allele. The bit-string model can indeed only be analysed with computer simulations. In contrast, for the Gaussian model we can analytically derive conditions under which to expect either a single generalist allele or the build-up of allelic diversity through gradual evolution in a process known as evolutionary branching (***Metz et al., 1992***; ***Geritz et al., 1998***; ***Kisdi and Geritz, 1999***; ***Doebeli, 2011***) (see ***SI Appendix 6 and 7*** for details).

**Figure 4.**
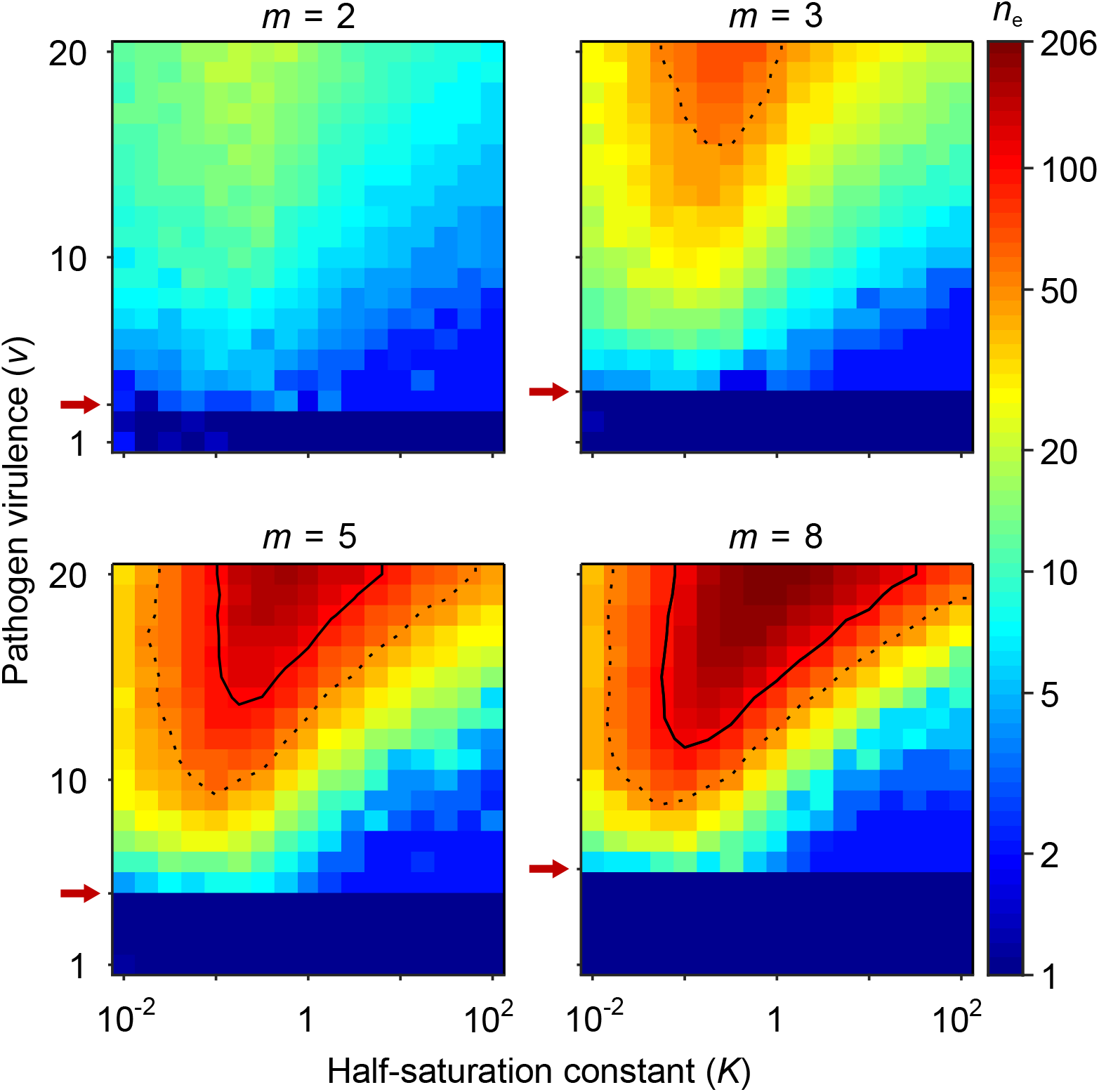
Number of coexisting alleles under the Gaussian model for *m* pathogens as a function of pathogen virulence *v* and the survival half-saturation constant *K*. Figures are based on a single individual-based simulation per pixel and run for 10^6^ generations, assuring that the equilibrium distribution of alleles is reached. Results are reported in terms of the effective number of alleles *n*_e_, which discounts for rare alleles present at mutation-drift balance (see ***Appendix 1***). The clear pattern in the figures indicates a high degree of determinism in the simulations. Results are reported in terms of the effective number of alleles *n*_e_, which is a conservative measure for the number of alleles, discounting for rare alleles present at mutation-drift balance. Using population size *N* = 10^5^ and per-capita mutation probability *µ* = 5 × 10^*−7*^, the expected *n*_e_ under mutation-drift balance alone equals 1.2 (see ***Appendix 1***). Dashed and solid lines give the contours for *n*_e_ = 50 and *n*_e_ = 100, respectively. Red arrows indicate 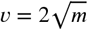.the threshold for polymorphism to emerge from branching (***Equation S46***). Accordingly, simulations in the dark blue area result in a single abundant allele with *n*_e_ close to one. Other parameters: expected mutational step size *δ* = 0.03.

## Results

### Gaussian Model

In the simulations of the Gaussian model, the evolutionary dynamics first proceed toward a generalist allele with an intermediate efficiency against all pathogens, to which we refer to as ***x***^*^. This generalist allele maximizes the condition *c* for homozygote genotypes (grey lines and cones in ***Figure 1, SI Appendix 7.2***). Once this generalist allele is reached, the evolutionary dynamics either stops (***Figure S1***), resulting in a population where all individuals are homozygous for ***x***^*^, or allelic diversification ensues (***Figure 3***), resulting in the coexistence of specialist and generalist alleles. Based on the adaptive dynamics approximation, we show analytically (***SI Appendix 7.1***) that ***x***^*^ is given by the arithmetic mean of the vectors ***p***_1_, …, ***p***_*m*_ describing the *m* pathogen optima (see ***Equation S27***) and an attractor of any sequence of allelic substitutions. Whether ***x***^*^ is an evolutionary stable endpoint or an evolutionary branching point where diversification ensues depends on the covariance matrix 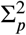 of the pathogen optima relative to the covariance matrices 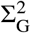 of the Gaussian efficiency functions (***SI Appendix 7.3-7.5)***For the special case of identically shaped Gaussian functions, diversification occurs if and only if

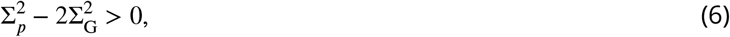

(***SI Appendix 7.4***). Note, that this expression is independent of the number of pathogens *m*. Under the additional assumption of equally distant pathogens and isotropic Gaussian functions, these covariance matrices are diagonal matrices with identical diagonal entries *σ*_*p*_ and *σ*_G_, respectively, and ***Condition 6*** simplifies to 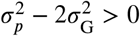. When pathogen optima having an equal distance of 1, the variance among the optima 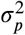 decreases with an increasing number of pathogens *m*, and the condition for evolutionary branching can be rewritten as

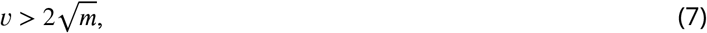

where 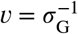 (***SI Appendix 7.5***).

***Figure 4*** presents the final number of coexisting alleles as derived from individual-based simulations. It shows that the number of coexisting alleles increases with the number of pathogens *m* and their virulence *v*, but also depends on the survival half-saturation constant ***K*** (***Equation 5***). For a large part of the parameter space, more than 100 (solid contour lines in ***Figure 4***) and up to over 200 alleles can emerge and coexist.

In order to better understand the process of allelic diversification, it is useful to inspect our analytical results in more detail. Evolutionary diversification occurs if mutant alleles *x*′ exist that can invade a population that is monomorphic for the generalist allele *x*^*^. Initially, while still rare, such mutant alleles will always occur in heterozygous individuals, where they are paired with the generalist allele. Thus, our condition for evolutionary diversification, 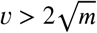, is equivalent to *s*(*x*′, *x*^*^) *> s*(*x*^*^, *x*^*^). Since, as homozygotes, the generalist allele maximizes condition and therefore survival (***SI Appendix 7.2***), we also have *s*(*x*^*^, *x*^*^) *> s*(*x*′, *x*′). In conclusion, individuals heterozygous for *x* conclusion, individuals heterozygous for and *x*^*^ have higher survival than either homozygote, *s*(*x*′, *x*^*^) *> s*(*x*^*^, *x*^*^) *> s*(*x*′, *x*′), and a polymorphism of these two alleles is maintained by HA, as suggested by ***Doherty and Zinkernagel*** (***1975***). Furthermore, the generalist allele is an evolutionary branching point in the sense of adaptive dynamics theory (***Geritz et al., 1998***; ***Kisdi and Geritz, 1999***).

The left-hand side of the diversification ***Condition 7***, indicates that invasion of more specialized alleles is favoured when pathogen virulence *v* is large (narrow Gaussian functions, see ***Figure 1A, C***). In this case, homozygotes for the generalist allele ***x***^*^ are relatively poorly protected against pathogens and more specialized alleles enjoy a fitness advantage while invading. The opposite is true when *v* is small (wide Gaussian functions, see ***Figure 1A, C***). The right-hand side of the diversification criterion indicates that the benefit of specialization decreases with an increasing number of pathogens (compare position of red arrows in ***Figure 4***), because different pathogens require different adaptations. Thus, counter to intuition, initial allelic diversification is disfavoured in the presence of many pathogens.

If initial allelic diversification occurs, it leads to a dimorphism from which new mutant alleles can invade if they are more specialized than the allele from which they originated. Then, two allelic lineages emerge from the generalist allele ***x***^*^ and subsequently diverge (***Figure 3A***, up to *t* = 3 × 10_4_ below grey plane). Increasing the difference between the two alleles present in such a dimorphism has two opposing effects. The condition and thereby the survival of the heterozygote genotype increases because the MHC molecules of the two more specialized alleles provide increasingly better protection against complementary sets of pathogens, that is, these alleles are subject to a divergent allele advantage (***Wakeland et al., 1990***; ***Pierini and Lenz, 2018***). On the other hand, survival of the two homozygote genotypes decreases because they become increasingly more vulnerable to the set of pathogens for which their MHC molecules do not offer protection. Note that, due to random mating and assuming equal allele frequencies, half of the population are high survival heterozygotes and the remaining half homozygotes with low survival. Since survival is a saturating function of condition *c* (***Equation 5***), it follows that the increase in survival of heterozygotes slows down with increasing condition (plateau of the orange curves in ***Figure 3***), and the two opposing forces eventually balance each other such that divergence comes to a halt. At this point, our simulations show that the allelic lineages can branch again, resulting in three coexisting alleles. As a result, the proportion of low-survival homozygotes decreases, assuming equal allele frequencies, from one-half to one-third. Subsequently, the coexisting alleles diverge further from each other because the increase in heterozygote survival once again outweighs the decreased survival of the (now less frequent) homozygotes (see ***Figure 3A***, at time *t* = 3 × 10^4^, grey plane). In ***Figure 3A***, this process of evolutionary branching and allelic divergence repeats itself one more time, resulting in four coexisting alleles. Consequently, ten genotypes emerge: four homozygotes and six heterozygotes. The homozygotes with specialist alleles have a condition, and thereby a survival, close to zero (two left bars in bottom panel). Conversely, the homozygote for the generalist allele ***x*** ^**\***^ has an intermediate condition (middle bar), and all heterozygote genotypes have a survival close to 1 (right bar).

In ***Figure 3B***-***D***, the process of evolutionary branching and allelic divergence continues to recur. As a consequence, allelic diversity continues to increase while simultaneously the proportion of vulnerable homozygote genotypes decreases (***Figure 3***, lower panel). Thus, in contrast to prior approaches (e.g. ***Kimura and Crow, 1964***; ***Wright, 1966***; ***Lewontin et al., 1978***; ***Maruyama and Nei, 1981***), we do not rely on hand-picked genotypic fitness values. Instead, in our approach, fitness values emerge from an eco-evolutionary model where evolution can be viewed as a self-organizing process finding large sets of alleles that can coexist (***Figure 3***, upper panel).

We note that the half-saturation constant ***K*** does not appear in the branching condition and thus does not affect whether polymorphism evolves. However, ***K*** does affect the final number of alleles, which is maximal for intermediate values of ***K***. This can be understood as follows. If ***K*** is very large (right-hand side of the panels in ***Figure 4***), then heterozygote survival saturates more slowly with increased allelic divergence so that continued allelic divergence is less counteracted. This hinders repeated branching (compare ***A*** and ***C*** in ***Figure 3***). On the other-hand, if ***K*** is very small (left-hand side of the panels in ***Figure 4***), then homozygous individuals can have high survival, which decreases the selective advantage of specialisation, leading to incomplete specialization and a reduced number of branching events (compare ***D*** and ***C*** in ***Figure 3***).

In summary, high virulence *v* promotes allelic diversification. Increasing the number of pathogens *m* has a dual effect: it hinders initial diversification but facilitates a higher number of coexisting alleles if diversification occurs, especially, for intermediate values of the half-saturation constant ***K***.

We perform several robustness checks. First, ***Figure S2*** shows simulations in which we vary the expected mutational step size. These simulations show that the gradual build-up of diversity occurs most readily as long as the mutational step size is neither very small, since then the evolutionary dynamics becomes exceedingly slow, nor very large, since a large fraction of the mutants are then deleterious and end up outside the simplex made up of the pathogen optima (e.g., outside the triangle made up by the three pathogen optima in ***Figure 1C-D***) so that they perform worse against all pathogens.

Second, the results presented in ***Figure 3*** and ***Figure 4*** are based on the assumptions of equally spaced pathogen optima and equal width and isotropic Gaussian functions *e*_*k*_(***x***) as shown in ***Figure 1***. In ***SI Appendix 7*** and ***SI Appendix 4***, we present analytical and simulation results, respectively, for the non-symmetric case. In particular, ***Figure S3*** shows that the predictions for the number of coexisting alleles presented here are qualitatively robust against deviations from symmetry. This is in line with ***Conditions 6*** and its simplification under full symmetry, 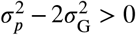.showing that the more general condition for the evolution of allelic polymorphism is structurally identical to the condition under full symmetry.

### Bit-String Model

Evolutionary diversification of MHC-alleles in the bit-string model is analysed with individual-based simulations, and the results are summarized in ***Figure 5***. Similar to the Gaussian model, we find high levels of allelic polymorphism, with over 100 alleles coexisting in a significant portion of the parameter space. Note that we here keep the half-saturation constant ***K*** fixed at 1. With this choice, the realized conditions occur both in the range where survival changes drastically with condition and where the survival function saturates (***Figure 6***), fulfilling assumption (b). This allows us to focus on the effect of the number of peptides *n*_pep_ per pathogen.

**Figure 5.**
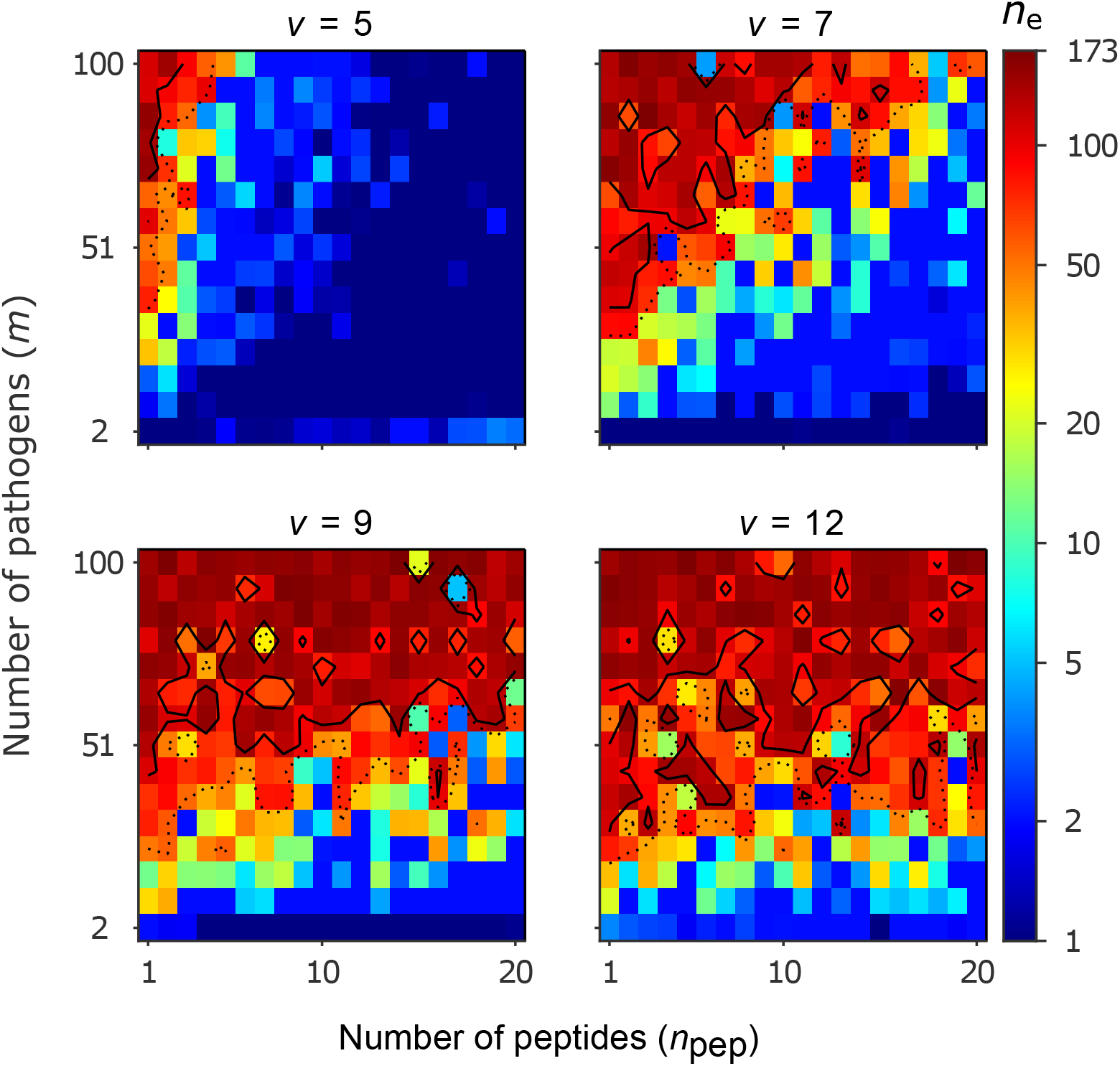
Number of coexisting alleles for the bit-string model for four values of virulence *v* as a function of the number of pathogens *m* (increased in steps of 7) and the number of peptides per pathogen *n*_pep_. Figures are based on a single individual-based simulation per pixel and run for 10^6^ generations. Results are reported in terms of the effective number of alleles *n*_e_, which discounts for rare alleles present at mutation-drift balance (see ***Appendix 1***). Using population size *N* = 10^5^ and per-capita mutation probability *µ* = 5 × 10^−6^, the expected *n*_e_ under mutation-drift balance alone equals 3. Dashed and solid lines give the contours for *n*_e_ = 50 and *n*_e_ = 100, respectively. Evolution started from populations monomorphic for a random allele, and run for 2 × 10^6^ generations, assuring that the equilibrium distribution of alleles is reached. Other parameters: half-saturation constant ***K*** = 1.

**Figure 6.**
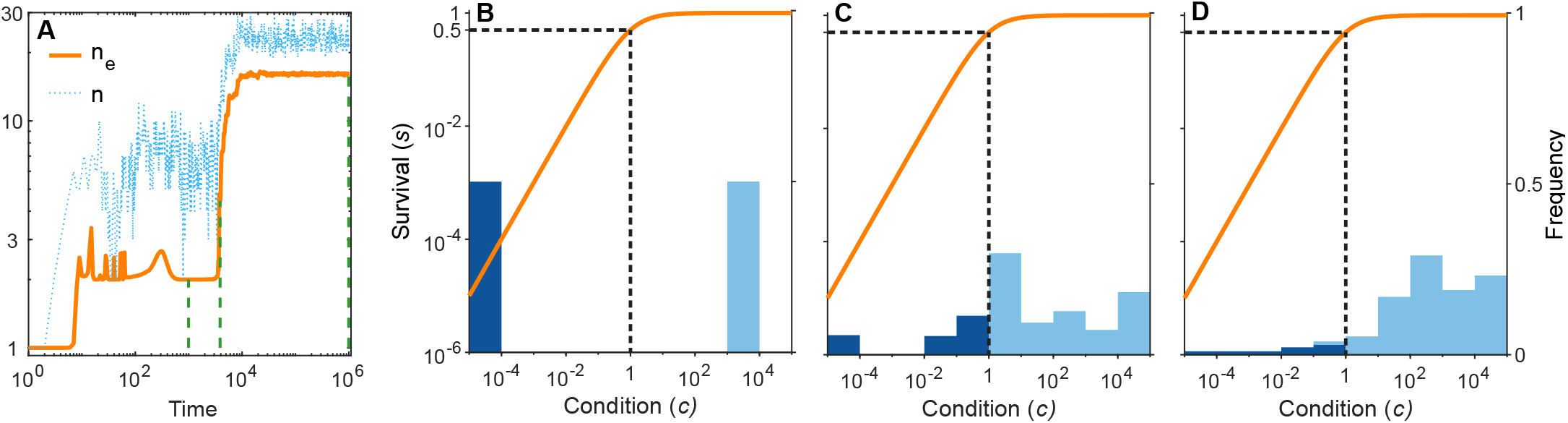
A simulation run showing the evolution of allelic diversity under the bit-string model in the presence of *m* = 12 pathogens. Panel ***A*** show the number of alleles *n* and the effective number of alleles *n*_e_ as a function of time (on a log-log scale). Panels ***B***-***D*** give survival *s* as a function of condition *c* as defined by ***Equation 5*** on a log-log scale (orange line, left vertical axis) and the distribution of conditions at three time points (***B***: *t* = 100, ***C***: *t* = 3900, ***D***: *t* = 10^6^; vertical green dashed lines in ***A***), with dark blue bars for homozygotes and light-blue bars for heterozygotes, right vertical axis (conditions from 0 to 10^−4^ are incorporated into the first bar, and conditions from 10^4^ and greater are incorporated in the last bar). This shows that as allelic diversity increases, the frequency of homozygotes with low survival decreases. The black dashed lines indicate the value of ***K*** = 1. Other parameter values: *v* = 7, *m* = 12, *n*_pep_ = 3, *N* = 10^5^, *µ* = 5 × 10^−6^.

Our results can be understood as follows. The likelihood that an MHC-molecule can recognize all pathogens is high in the following region of the parameter space. Firstly, if virulence *v* is low, then peptide recognition is more likely (***Equation 2***). Secondly, if the number of pathogens *m* is low, then detection of all pathogens is a simpler task. Thirdly, if the number of peptides *n*_pep_ per pathogen is high, then the potential for successful pathogen detection increases (***Equation 3***). Although our model is not sufficiently mechanistic to be directly related to parameters observed in nature, it suggests that when pathogens have a high number of peptides, maintaining allelic polymorphism requires a larger number of pathogens under conditions of low virulence (*v* ≤ 7). For higher virulence (*v* ≥ 9), the effect of *n*_pep_ weakens, and allelic polymorphism evolves seemingly independent of the number of pathogens. Each of these three circumstances facilitates the existence of a single best allele whose MHC molecule recognizes all pathogens with a high probability (dark blue regions in ***Figure 5***).

As virulence or the number of pathogens increases, or as the number of peptides decreases, the task of recognizing all pathogens with high probability becomes progressively more challenging. This leaves homozygous individuals vulnerable to an increasing array of pathogens. As homozygotes get more vulnerable, there is a growing advantage for heterozygotes carrying alleles with complementary immune profiles, as these are able to detect up to twice as many pathogens as either homozygote. This increasingly stronger HA, in turn, facilitates coexistence of an increasing number of alleles, illustrated by increasingly warmer colours in ***Figure 5***, and thereby decreases the proportion of vulnerable homozygotes. Thus, similar to the Gaussian model, increasing either the virulence *v* or the number of pathogens *m* enables a higher number of alleles to coexist. However, unlike the Gaussian model, increasing *m* actually facilitates initial diversification rather than hindering it.

Importantly, in the bit-string model, a point mutation, switching the value of an arbitrary bit in the bit-string, can drastically alter the efficiencies against a large set of pathogens. Because of this, and in contrast to the Gaussian model, a polymorphism maintained by HA can emerge from many different alleles. On the other hand, gradual evolution in the Gaussian model is more efficient in finding the evolutionary end-point of complementary alleles (***Figure 3***), while for the bit-string model, as the number of alleles increases, this becomes slower due to the lack of fine-tuning as mutations have large effect. To compensate for this, we use, compared to the Gaussian model, a higher mutation probability *µ* and run simulations for more generations.

***Figure 6A*** shows the build-up of allelic diversity over time in an exemplary simulation run, and ***B***-***D*** show the distribution of condition values at three time points, as indicated by green hatched lines in ***A***. In ***B*** the population is dimorphic. Due to random mating, half of the population consists of homozygotes with low condition (dark blue bar), while the remaining half are heterozygotes with high condition (light blue bar). As time proceeds, the number of coexisting alleles increases. ***Figure 6C*** depicts a stage with five coexisting alleles (with at least 1% frequency) and an effective number of alleles (*n*_e_) of 4.4. Ultimately, evolution results in 19 coexisting alleles (with at least 1% frequency), and an *n*_e_ of 16.1, as shown in ***D***. In this process, the number of low-condition homozygotes decreases, as indicated by the dark blue bars.

## Discussion

Heterozygote advantage (HA) as an explanation for the coexistence of a large number of alleles at a single locus has a long history in evolutionary genetics. ***Kimura and Crow*** (***1964***) and subsequently ***Wright*** (***1966***) showed that HA can in principle result in the coexistence of an arbitrary number of alleles at a single locus if two conditions are met: (1) all heterozygotes have a similarly high fitness, and (2) all homozygotes have a similarly low fitness. One special class of genes fulfilling these assumptions occur at self-incompatibility loci, where mating partners need to carry different alleles for fertilisation to be successful (***Wright, 1939***; ***Castric and Vekemans, 2004***), or loci where homozygosity is lethal (***Ding et al., 2021***). However, more generally these conditions were deemed unrealistic by Kimura, Crow and Wright themselves. This assessment was subsequently confirmed by ***Lewontin et al. (1978***), who investigated a model in which the exact fitnesses are determined by drawing random numbers in a manner that all heterozygotes are more fit than all homozygotes. They found that the proportion of fitness arrays that leads to a stable equilibrium of more than six or seven alleles is vanishingly small. Similarly, the idea that the high allelic diversity found at MHC loci can be explained by HA was initially accepted by theoreticians (e.g. ***Maruyama and Nei, 1981***; ***Takahata and Nei, 1990***), but several later authors studying models based on more mechanistic assumptions were unable to reliably predict the coexistence of significantly more than ten alleles (***Spencer and Marks, 1988***; ***Hedrick, 2002***; ***de Boer et al., 2004***; ***Borghans et al., 2004***; ***Stoffels and Spencer, 2008***; ***Trotter and Spencer, 2008, 2013***; ***Ejsmond and Radwan, 2015***; ***Lau et al., 2015***). Thus, currently HA is largely dismissed as an explanation for highly polymorphic loci (***Gould et al., 2004***; ***Eizaguirre and Lenz, 2010***; ***Lenz, 2011***; ***Hedrick, 2012***).

Our study, while not meant to be a highly realistic mechanistic representations of the interaction between MHC genes and pathogens, serves as a proof of principle that a high number of alleles, matching those found at MHC loci in natural populations, can indeed arise in an evolutionary process driven by HA. Our results thus revive the idea that HA has the potential to explain extraordinary allelic diversity. Importantly, and in contrast to several of the above-mentioned studies, this is achieved without making direct assumptions about homozygote and heterozygote fitnesses. Instead, our results emerge from two assumptions about how pathogens affect a host’s condition and how this, in turn, affect survival. Assumption (a) states that pathogens are lethal in the absence of an appropriate immune response. This assumption is implemented in our model by assuming that each pathogen decreases a host’s condition in a proportional manner (***Equation 4***), rather than by a fixed amount. Assumption (b) states that the effect of pathogens depends on host condition, with hosts in poorer condition being affected more strongly. Then, the combined effect of multiple pathogens on host survival exceeds the sum of the effects of each pathogen alone. Thus, many pathogens against which a host has an imperfect immune response can collectively push a host’s condition below a threshold where mortality becomes rather high (orange lines in ***Figure 3*** and ***Figure 6***). In our model, this assumption is fulfilled rather naturally. Since the probability to survive can logically not exceed 1, the function that maps condition to survival has to be saturating (***Equation 5***).

In the following, we detail how assumptions (a) and (b) can result in the emergence of well over 100 alleles such that heterozygotes have similarly high fitness (condition (1) of ***Kimura and Crow***) and homozygotes have similarly low fitness (condition (2) of ***Kimura and Crow***). We start with the observation that the survival probabilities in evolved polymorphic populations vary between individuals (lower panels in ***Figure 4*** and ***Figure 5B-D***). Part of the population consists of individuals that have very low survival probabilities. These are individuals with a condition value considerably less than ***K*** and they are almost exclusively homozygotes. This is because, whenever polymorphism is favoured, homozygotes are poorly defended against some pathogens and the fact that pathogens affect condition multiplicatively (***Equation 4***). The remaining part of the population consists of individuals with condition values considerably above ***K***. Although the condition of these individuals can differ by several orders of magnitude, their survival is close to 1, which results from the fact that the function that maps condition to survival is saturating. These individuals are almost exclusively heterozygotes. This is because alleles that protect against complementary sets of pathogens, when paired together, offer at least a decent protection against all pathogens. In summary, our assumptions (a) and (b) lead to a set of alleles such that their survival probabilities fall into two clusters as required for condition (1) and (2) of ***Kimura and Crow*** (***1964***) to be fulfilled. The larger the number of alleles, the lower becomes the proportion of vulnerable homozygotes, and the population consists increasingly of almost equally fit heterozygotes.

***Borghans et al. (2004***) use a bit-string model similar to ours with *m* = 50 pathogens, *n*_pep_ = 20 peptides, a virulence of *v* = 7 and a step function for the probability that an MHC molecule detects a peptide (*a* → ∞ in ***Equation 2***). In contrast to our model, they assume that an individual’s condition equals the proportion of detected pathogens, meaning that each pathogen can reduce fitness by only 2% (thereby not fulfilling our assumption a). Additionally, they assume that survival is proportional to the squared condition (not fulfilling our assumption b). ***Figure S4*** in ***SI Appendix 5*** shows a run of our bit-string model with the parameter values used by ***Borghans et al. (2004***), resulting in more than 100 coexisting alleles. In contrast, they find only up to seven coexisting alleles, demonstrating that assumption (a) and (b) in our model drive the high number of coexisting alleles found by us.

Currently, there are several mechanisms proposed to explain the diversity observed at MHC loci. First, in the presence of an HA, each allele has an advantage when rare because it almost always occurs in heterozygotes. Thus, there is negative frequency-dependent selection acting at the level of the allele. In addition, negative frequency-dependent selection can arise from, for example, Red-Queen dynamics, fluctuating selection and disassortative mating (***Apanius et al., 1997***; ***Hedrick, 1999***; ***Penn, 2002***; ***Borghans et al., 2004***; ***Wegner, 2008***; ***Spurgin and Richardson, 2010***; ***Loiseau et al., 2011***; ***Ejsmond and Radwan, 2015***; ***Lighten et al., 2017***; ***Ejsmond et al., 2023***). These mechanisms are similar to HA in the sense that the selective advantage of an allele increases with decreasing frequency. However, they do not result in heterozygotes being more fit than the homozygotes carrying the rare allele. In addition, neutral diversity can be enhanced by recombination (***Klitz et al., 2012***; ***Linnenbrink et al., 2018***; ***Robinson et al., 2017***). If many individuals are heterozygous, the particularly high levels of gene conversion found at MHC genes can be effective in creating new allelic variants. For instance, for urban human populations with a large effective population size of *N*_e_ = 10^6^ and a per-capita gene conversion probability of *r* = 10^−4^ an effective number of alleles as high as *n*_e_ = 1 + 4*rN*_e_ = 401 can theoretically be maintained by gene conversion (***Klitz et al., 2012***). However, it is important to point out that for gene conversion to increase allelic diversity, some genetic polymorphism due to balancing selection has to exist to start with. We do not claim that the mechanisms listed here do not play an important role in maintaining allelic diversity at MHC loci. Rather, our results show that, contrary to the currently widespread view, HA should not be dismissed as a potent force. In any real system, different mechanisms will jointly affect allelic diversity. For instance, ***Lighten et al. (2017***) present a model in which, for Red-Queen co-evolution to maintain allelic polymorphism, HA in the form of a divergent allele advantage (***Wakeland et al., 1990***) seems to be a necessary ingredient. Similarly, ***Borghans et al. (2004***) show that pathogen co-evolution can further increase the number of co-existing alleles compared to HA alone.

The aim of our study is to understand how HA on its own can result in allelic polymorphism. For this reason, we kept all aspects concerning pathogens fixed, focusing on a scenario where pathogen optima represent diverse taxonomic groups that remain approximately constant over the time scales considered in our model. This approach excludes Red-Queen dynamics and fluctuating selection. Models of Red-Queen dynamics are based that pathogens evolve to avoid detection by the host’s immune system (***Borghans et al., 2004***; ***Ejsmond and Radwan, 2015***; ***Ejsmond et al., 2023***). In our model, this would correspond to moving pathogen optima (in the Gaussian model) or changes in the pathogen peptides (in the bit-string model). We expect that incorporating this would hamper the build-up of allelic MHC diversity when driven solely by HA if pathogens evolve quickly. Alleles previously maintained as beneficial would then become disadvantageous and go extinct more rapidly than new advantageous alleles can appear.

Another component of pathogens that can evolve in response to host immune defence is their virulence (***Frank and Schmid-Hempel, 2008***). The transmission-virulence trade-off hypothesis (***Anderson and May, 1982***; ***Frank, 1996***; ***Alizon et al., 2009***) predicts that pathogens that cause relatively little harm to their host (i.e., pathogens with low virulence) may evolve towards higher virulence to increase their transmission rate. In line with this hypothesis, we speculate that incorporating virulence evolution leads to higher virulence whenever pathogens inflict little harm on their hosts. This scenario applies in the dark blue parameter regions in ***Figure 4*** and ***Figure 5***, where host populations possess a single effective generalist allele. In these regions, the evolution of increased virulence would shift pathogens into parameter regions where allelic polymorphism becomes adaptive. The ensuing build-up of allelic polymorphism decreases the harm inflicted by pathogens through HA, which, in turn, increases the selection pressure acting on pathogens for an even further increase in virulence. This suggests, in contrast to evolving pathogen optima, a positive feedback-loop between virulence evolution and the evolution of allelic diversity.

Our Gaussian model is not restricted to MHC genes, but can apply to any gene that affects several functions important for survival. Examples are genes that are expressed in different ontogenetic stages or different tissues with competing demands on the optimal gene product. However, gene duplication is expected to reduce the potential number of coexisting alleles per locus and eventually lead to a situation where the number of duplicates equals the number of functions (***Proulx and Phillips, 2006***). Under this scenario, the high degree of polymorphism reported here would be transient. However, for MHC genes evidence exist that other forces limit the number of MHC loci (***Penn, 2002***; ***Wegner, 2008***; ***Eizaguirre and Lenz, 2010***; ***Spurgin and Richardson, 2010***). But it is important to point out that, while our model focuses on evolution at a single MHC locus, many vertebrates have more than one MHC locus with similar functions (***Wegner, 2008***; ***Eizaguirre and Lenz, 2010***; ***Spurgin and Richardson, 2010***). The diversity generating mechanism described here still applies if the different loci are responsible for largely non-overlapping sets of pathogens, indicating that the mechanism presented here can in principle explain the high number of coexisting MHC alleles.

In summary, our research offers a fresh view that can help us to understand allelic diversity at MHC loci. We identify two crucial assumptions related to pathogen-host interactions, under which we show that heterozygote advantage emerges as a potent force capable of driving the evolution of a very high number of coexisting alleles.

## Appendix 1

Here, we provide the calculations for the effective number of alleles *n*_e_ reported in *Figures 4* and *5*. The effective number of alleles is given by the reciprocal of the population homozygosity 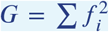.where *f*_i_ denotes the frequency of allele *i* in the population (Kimura and Crow, 1964). Under mutation-drift balance, the expected homozygosity is approximated by 1/(1 + 4*N μ*) (Gillespie, 2004), where *N* is population size and *μ*) the per-capita mutation probability.

Thus, under mutation-drift balance, the expected value of *n*_e_ equals 1+4 *N μ*. For *Figure 4*, where *N* = 10^5^ and *μ* = 5 × 10^− 7^, the expected value of *n*_*e*_ is 1.2. In *Figure 5*, with *N* = 10^5^ and *μ* = 5 × 10 ^− 6^, the expected value for *n*_e_ is 3. Hence, *n*_*e*_-values significantly higher than these expectations indicate the presence of alleles maintained by balancing selection.

It is worth noting that when alleles are at equal frequencies *f*_i_ = 1/*n, n*_*e*_ is equal to n. In our model, both condition (1) and (2) of Kimura and Crow (1964) are approached at evolutionary equilibrium (i.e., heterozygote having similar and high fitness while homozygote having similar and low fitness), as elaborated in the Discussion. As a result, alleles maintained by HA are maintained at roughly similar frequencies. Consequently, *n*_*e*_ gives a good estimate for the number of alleles that coexist in a protected polymorphism due to HA, rather than being maintained in a balance between mutation and drift.

## Author Contributions

Conceptualization: M.S. and C.R.; Methodology: M.S. and C.R.; Formal analysis: M.S.; Visualization: M.S.; Writing: M.S. and C.R.; Supervision: C.R.; Funding Acquisition: C.R.

## Acknowledgements

We thank Yvonne Meyer-Lucht and Tobias Lenz for helpful discussions and Göran Arnqvist, Helena Westerdahl, Joachim Hermisson and Sophie Karrenberg for comments on an earlier version of the manuscript.

## Supplementary Information

### 1. Supplementary Figure S1: Evolutionary Dynamics Without Diversification

**Fig. S1.**
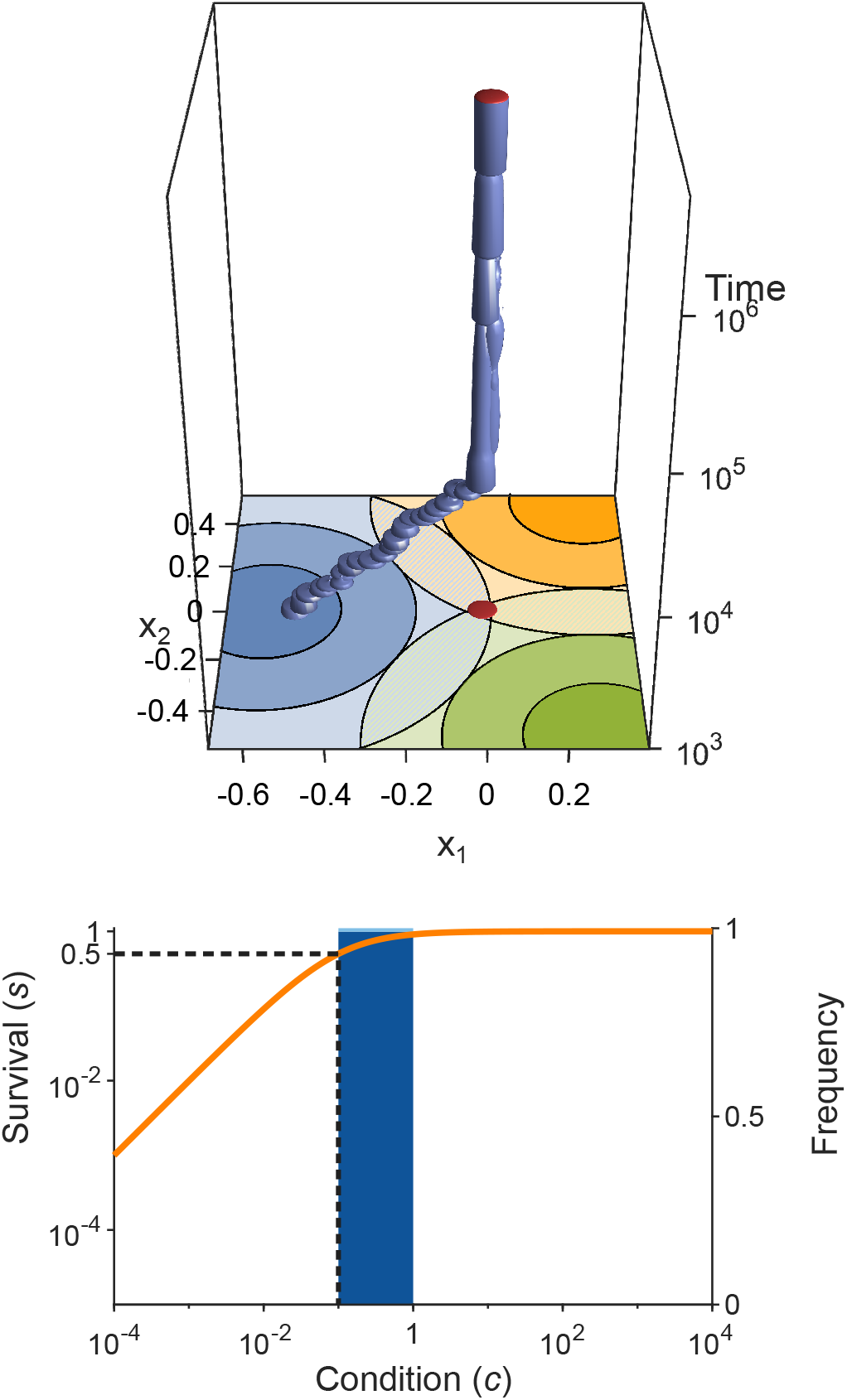
Evolution of allelic values in the presence of three pathogens. This figure is analogous to ***Figure 3*** (see that legend for details) but with wider Gaussians (*v* = 2.5, as in ***Figure 1C***). As a consequence, the condition for evolutionary branching 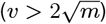is not fulfilled and the evolutionary dynamics result in a monomorphic population consisting essentially of only the generalist allele ***x***^*^ = (0, 0). This result is independent of the half-saturation constant *K*, here chosen to be *K* = 10.

### 2. Table of Mathematical Notation

List of all mathematical symbols used in the Supplementary Information. Bold italic font indicates vectors (e.g., ***x***) while normal italic font indicates numbers or scalar-valued functions. Capital letters in sans serif font indicate matrices (e.g., H).

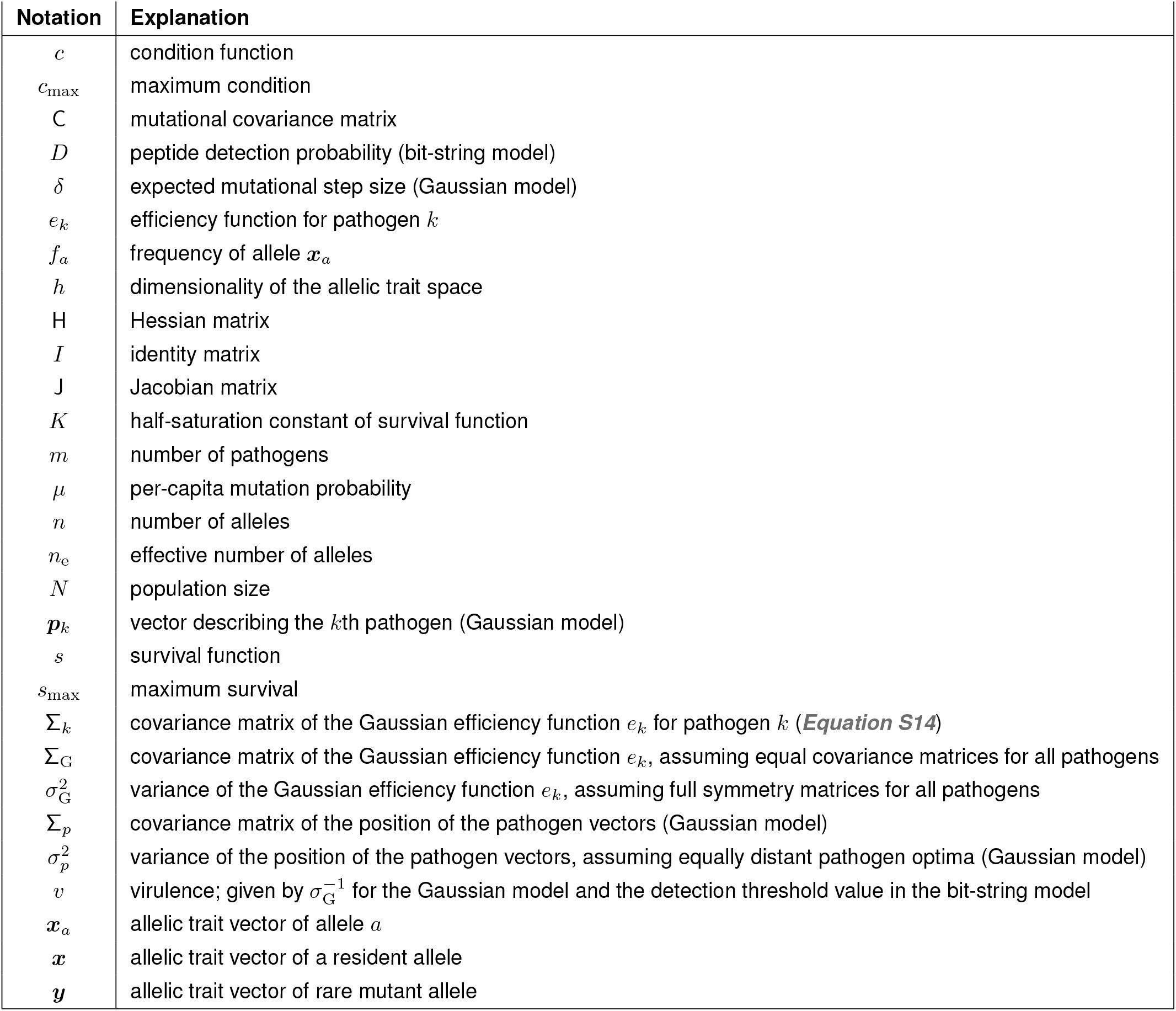

| Notation | Explanation |
| --- | --- |
| $c$ | condition function |
| $c_{\max}$ | maximum condition |
| $\mathbf{C}$ | mutational covariance matrix |
| $D$ | peptide detection probability (bit-string model) |
| $\delta$ | expected mutational step size (Gaussian model) |
| $e_k$ | efficiency function for pathogen $k$ |
| $f_a$ | frequency of allele $x_a$ |
| $h$ | dimensionality of the allelic trait space |
| $\mathbf{H}$ | Hessian matrix |
| $\mathbf{I}$ | identity matrix |
| $\mathbf{J}$ | Jacobian matrix |
| $K$ | half-saturation constant of survival function |
| $m$ | number of pathogens |
| $\mu$ | per-capita mutation probability |
| $n$ | number of alleles |
| $n_e$ | effective number of alleles |
| $N$ | population size |
| $\mathbf{p}_k$ | vector describing the $k$th pathogen (Gaussian model) |
| $s$ | survival function |
| $s_{\max}$ | maximum survival |
| $\Sigma_k$ | covariance matrix of the Gaussian efficiency function $e_k$ for pathogen $k$ (**Equation S14**) |
| $\Sigma_G$ | covariance matrix of the Gaussian efficiency function $e_k$, assuming equal covariance matrices for all pathogens |
| $\sigma_G^2$ | variance of the Gaussian efficiency function $e_k$, assuming full symmetry matrices for all pathogens |
| $\Sigma_p$ | covariance matrix of the position of the pathogen vectors (Gaussian model) |
| $\sigma_p^2$ | variance of the position of the pathogen vectors, assuming equally distant pathogen optima (Gaussian model) |
| $v$ | virulence; given by $\sigma_G^{-1}$ for the Gaussian model and the detection threshold value in the bit-string model |
| $\mathbf{x}_a$ | allelic trait vector of allele $a$ |
| $\mathbf{x}$ | allelic trait vector of a resident allele |
| $\mathbf{y}$ | allelic trait vector of rare mutant allele |

### 3. Varying the Expected Mutational Step Size in the Gaussian Model

For the Gaussian model, mutations are drawn from an isotropic normal distribution, i.e., a matrix with covariance matrix *σ*_*µ*_*I* of dimension *h*. The expected mutational step-size *δ* is given by *σ*_*µ*_ times the expected value of the Chi-distribution (Equation 18.14 in ***Johnson et al., 1994***),

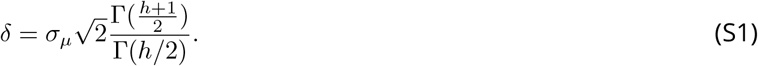

**Fig. S2.**
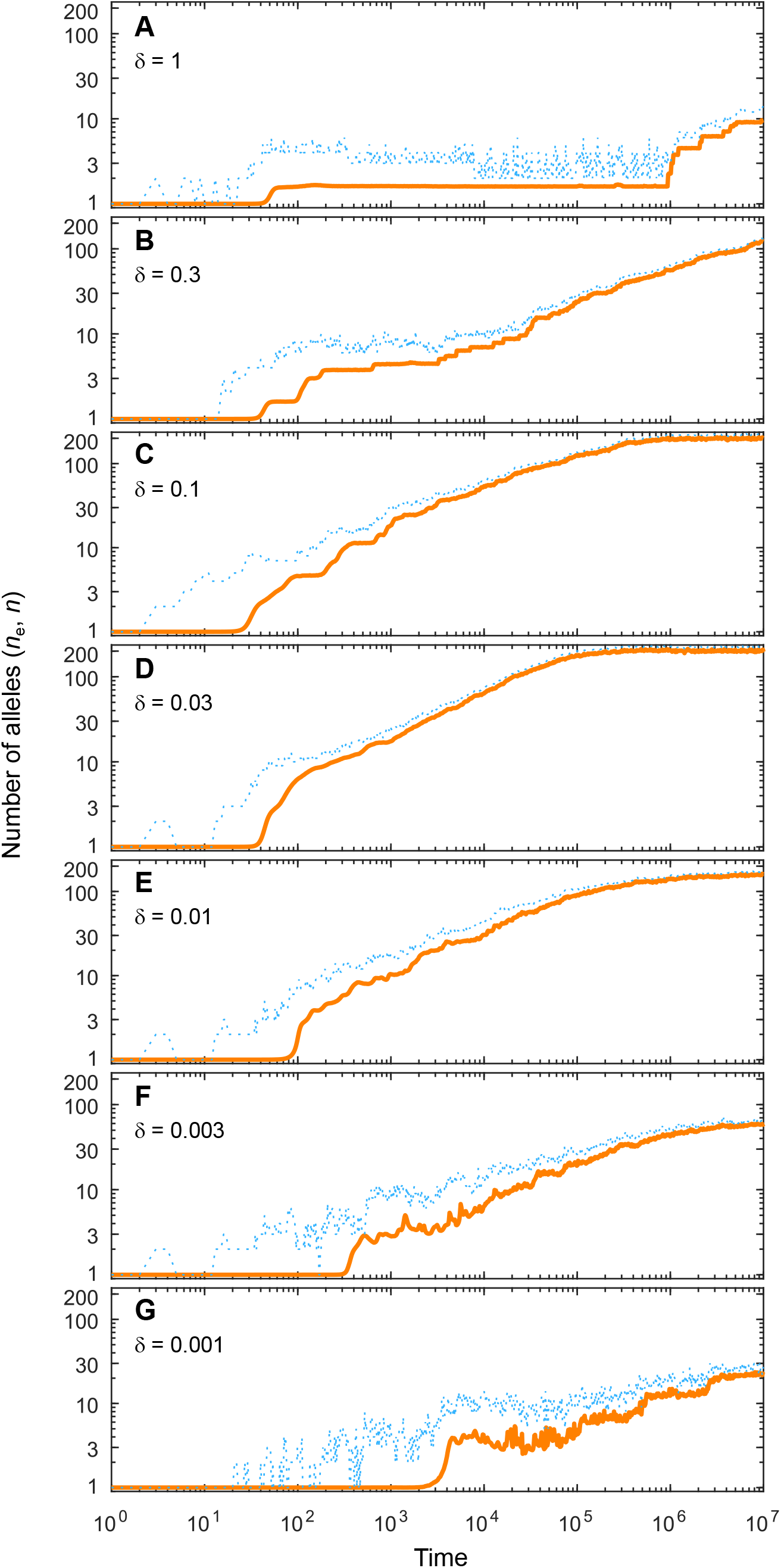
Number of coexisting alleles as they emerge in individual-based simulations for different expected mutational step sizes *δ* and eight pathogens (*m* = 8). Parameters are chosen such that up to 200 alleles can evolve (*K* = 0.5, *v* = 20; see bottom right panel in ***Figure 4*** in the main text). Solid orange lines and dotted blue lines give the effective number *n*_e_ and the absolute number *n* of alleles, respectively. The number of alleles increases fastest and saturates earliest for an intermediate expected mutational step size of *δ* = 0.03 (***D***; pathogen vectors are 1*/δ* = 1*/*0.03 ≈ 30 average mutational steps apart) as used in ***Figure 4***. Decreasing the average mutational step size slows down the build-up of allelic diversity (***E*** -***G***). In the extreme case shown in ***G*** (pathogen vectors are 1*/*0.001 = 1000 average mutational steps apart), the evolutionary dynamics is strongly limited by the rate of phenotypic change due to the small step size and the number of alleles after 10^7^ time steps has reached only 10% the number reached in ***D***. Increasing the average mutational step size also slows down the build-up of allelic diversity (***A***-***C***). In the extreme case shown in ***A*** (pathogen vectors are 1.25 average mutational steps apart), the evolutionary dynamics are strongly limited by the very large proportion of maladapted mutants. Other parameters (as in ***Figure 4***): *N* = 10^5^, *µ* = 5 *×* 10^−7^.

### 4. Deviations from Symmetry in the Gaussian Model

The number of coexisting alleles for different parameter combinations are shown in ***Figure 4*** in the main text. These results are based on two symmetry assumptions. First, the *m* points describing by the pathogen vectors are placed equidistantly with *d* = 1, resulting in a regular (*m* − 1)-simplex. Second, the multivariate Gaussian functions *e*_*k*_ describing the MHC-molecule’s efficiencies against the different pathogens are isotropic and have equal width, as shown in ***Figure 1***. Thus, the covariance matrices Σ_*k*_ in ***Equation S14*** are equal to 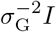.where *I* is the identity matrix. Here, we test the robustness of the outcome shown ***Figure 4*** with respect to violations of these symmetry assumptions. We focus on the case with eight pathogens, and the results are summarized in ***Figure S3***. Panel (***A***) is identical to the bottom right panel in ***Figure 4***, and shown here for comparison. Panels (***B-D***) show the final number of coexisting alleles for increasing deviations from symmetry, as explained in the following. Note that each pixel in the figure is based on a single simulation with a unique random perturbation from symmetry.

In ***Figure S3B*** the assumption of symmetrically placed pathogen vectors is perturbed while the Gaussian functions *e*_*k*_ are kept rotationally symmetric with equal width. ***Section 4.1*** describes the procedure how the positions of the pathogen vectors are randomized. The similarity between panel (***A***) and (***B***) indicates that deviations from a symmetric placement of pathogen vectors has a minor effect on the number of coexisting alleles. The slightly decreased smoothness of the contours corresponding to 50 and 100 coexisting alleles stems from the fact that each simulation (corresponding to a pixel) is based from a unique perturbation. Note that polymorphism can emerge for values of *v* such that the branching condition 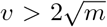derived for the symmetric case is not fulfilled (below the red arrow). This can be understood based on the expression for the Hessian matrix given in ***Equation S43***. This Hessian matrix is more likely to be positive definite for asymmetrically placed pathogen vectors.

In ***Figure S3C*** and ***D*** we, additionally to the non-symmetric placement of pathogen vectors, allow for Gaussian functions *e*_*k*_ that are not isotropic. The variances of the perturbed covariance matrices are drawn from the interval 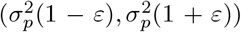 and constrained such that the average variance is equals 1, and then rotated randomly. ***Section 4.2*** describes this procedure in detail. Panel (***C***) shows the result for modest (*ε* = 0.2) and panel (***D***) for strong (*ε* = 0.5) deviations from rotational symmetry. Comparing panel (***C***) to (***B***) indicates that modest deviations from rotational symmetry have a relatively minor effect on the final number of coexisting alleles. In contrast, in panel (***D***) configurations exist where significantly fewer alleles are able to coexist. Interestingly, configurations resulting in a high number of alleles are more likely to occur in combination with high *K*-values. The highly irregular pattern results from each pixel corresponding to a single simulation with a unique random perturbation from symmetry. Furthermore, the threshold for polymorphism decreases even more because the Hessian matrix given in ***Equation S9*** is even more likely to be positive definite with perturbations in Σ_*k*_.

#### 4.1 Random Placement of Pathogen Vectors

We here describe how we randomly place eight pathogen vectors in trait space. In order to keep the results comparable to the symmetric case, we keep the average variance calculated from the position of their mid-points constant. The distribution of eight pathogen vectors can be described by their seven dimensional covariance matrix Σ_*p*_ calculated from the coordinates ***p***_1_, …, ***p***_8_. Since each diagonal element of Σ_*p*_ describes the variance of the pathogen vectors along a different dimension of the trait space, the average variance equals tr(Σ_*p*_)*/*7, where tr(Σ_*p*_) denotes the trace. This measure is unaffected by rotation of the points ***p***_1_, …, ***p***_8_. For symmetrically placed pathogen vectors Σ_p,sym_ = *d*^2^*/*(2*m*)*I*, where *I* denotes the identity matrix, and therefore tr(Σ_p,sym_) = *d*^2^(*m* − 1)*/*(2*m*). For the pathogens with perturbed placements (with covariance matrix Σ_p,per_), we demand tr(Σ_p,per_) = tr(Σ_p,sym_). We achieve this by first choosing eight preliminary points 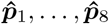 that are placed randomly within a unit 7-sphere, having the covariance matrix 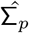. By subsequently multiplying the coordinates of these points by a scalar *α*, the variances in 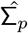 are multiplied by *α*^2^. By setting

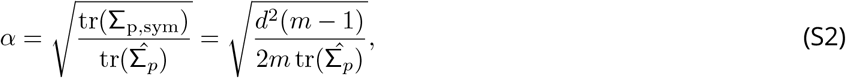

with *m* = 8, we obtain the final set of pathogen vectors ***p***_1_, …, ***p***_8_ with a covariance matrix Σ_p,per_ fulfilling tr(Σ_p,per_) = tr(Σ_p,sym_).

#### 4.2 Random Covariance Matrices for the Pathogen Efficiencies

We here describe how we create random covariance matrices Σ_*k*_. In order to keep the results between the symmetric and asymmetric case comparable, we fix the mean variance over all Σ_*k*_ to *σ*^2^ = *v*^−2^. We obtain the eight random covariance matrices Σ_1_, …, Σ_8_ in the following manner. First, eight random diagonal matrices D_1_, …, D_8_ are determined (one per pathogen vectors) with entries drawn from a uniform distribution *U*(1 − *ε*, 1 + *ε*). These matrices are then multiplied with the scalar

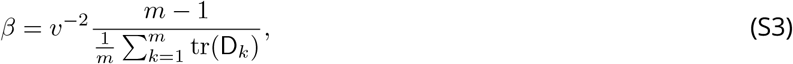

with *m* = 8 to obtain the set of matrices M_1_, …, M_8_ obeying 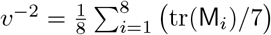.In a final step, we draw eight random rotation matrices R_1_, …, R_8_ and calculate our final covariance matrices P_1_, …, P_8_ as 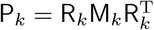.

**Fig. S3.**
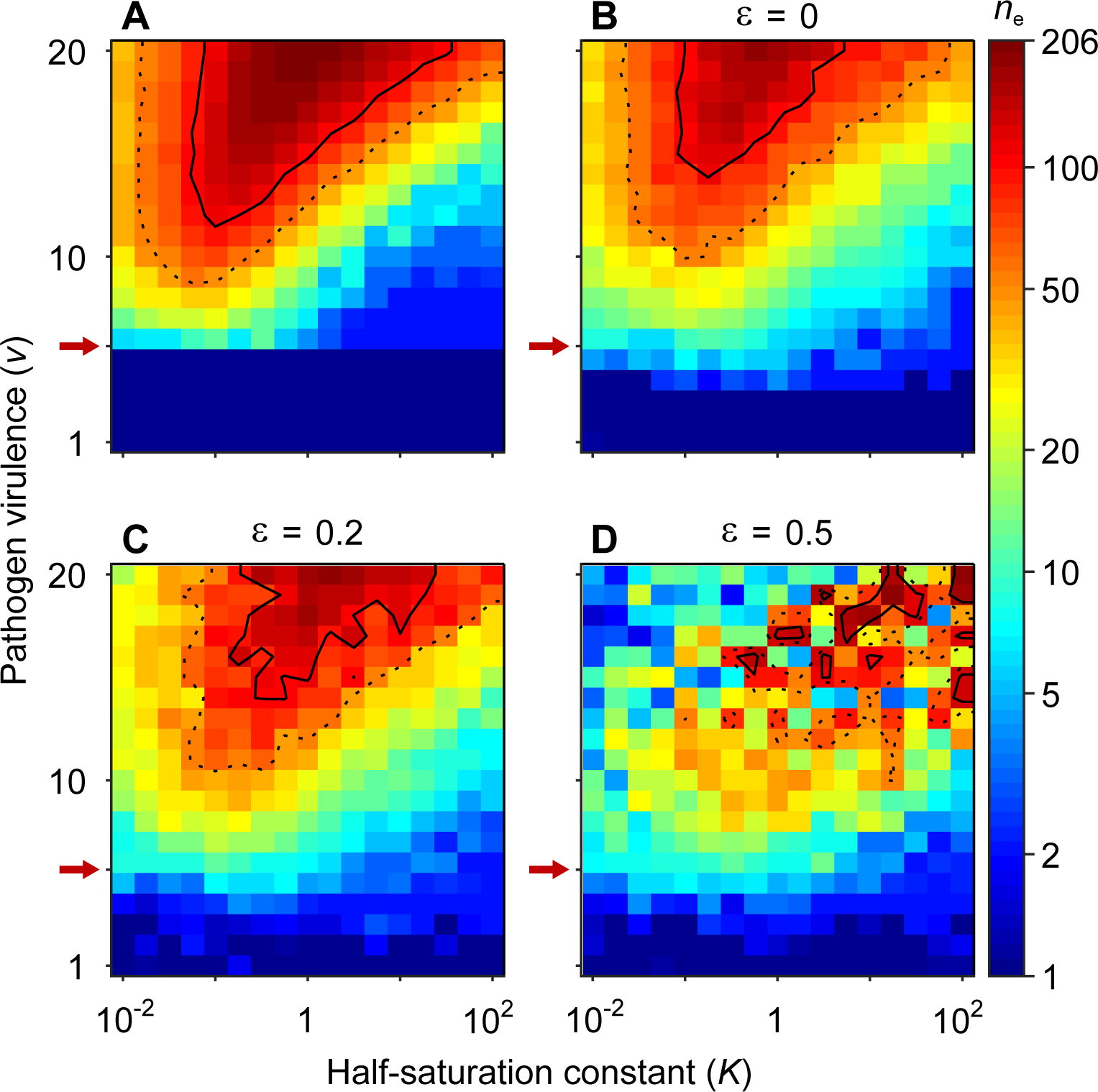
Number of coexisting alleles for eight pathogens as a function of pathogen virulence *v* and the half-saturation constant *K* for symmetrically (***A***) and non-symmetrically placed pathogen vectors (***B-D***). Figures are based on a single individual-based simulation per pixel and run for 10^6^ generations, assuring that the equilibrium distribution of alleles is reached. Panel (***A***) shows results for equally spaced pathogen vectors and isotropic functions *e*_*k*_ (***Equation S14***). It is identical to the bottom right panel in ***Figure 4*** and shown here for comparison. Panel (***B-D***) show the result for increasing perturbations from symmetry. In panel (***B***), pathogen vectors are placed randomly (see ***Section 4.1*** for details) while the functions *e*_*k*_ are kept rotationally symmetric. In panel (***C***) and (***D***), additionally to the non-symmetric placement of pathogen vectors, the functions *e*_*k*_ are independently perturbed from rotational symmetry (see Appendix 4.2 for details). In panel (***C***) the deviations from rotational symmetry are moderate, while in panel (***D***) they are strong. Note that in panel (***B-D***) pathogen vectors are no longer at a constant distance 1, but instead have the mean variance calculated from the pathogen optima corresponds to the variance of symmetrically placed pathogens optima with distance 1. Results are reported in terms of the effective number of alleles *n*_e_, which discounts for alleles arising from mutation-drift balance (see ***Appendix 1***). Dashed and solid lines give the contours for *n*_e_ = 50 and *n*_e_ = 100, respectively. Red arrows indicate 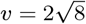.the minimal value for polymorphism to emerge from branching under full symmetry (***Equation S46***). Accordingly, simulations in the dark blue area result in a single abundant allele with *n*_*e*_ close to one. Other parameters: population size *N* = 10^5^, per-capita mutation probability *µ* = 5 *×* 10^−7^, expected mutational step size *δ* = 0.03.

### 5. Simulation Run of the Bit-String Model with Parameter Values as in Borghans *et al*. (2004)

We here present a comparison of our bit-string model with that of Borghans et al. (2004). These authors analyse a bit-string model with *m* = 50 pathogens (that are allowed to mutate but are not subject to selection), with *n*_pep_ = 20 peptides each, a virulence of *v* = 7 (with a step function for the probability that an MHC molecule detects a peptide), a population size of *N* = 10^3^ and a per-capita mutation probability of *µ* = 10^−5^. In contrast to our model, they assume that condition equals the proportion of detected pathogens, such that each pathogen can lower fitness by only 2% (not fulfilling our assumption a) and that survival is proportional to the squared condition (not fulfilling our assumption b). With these parameters and parameter values, their simulation results in up to seven alleles. We note, that the effective number of alleles in these simulations is likely lower, but no allele frequencies are given.

We contrast their results with those from our model, which, as detailed in the main part, fulfils assumptions a) and b). To approximate the step function for the detection probability, we use

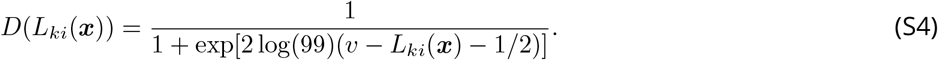

For this function, *v* is the required match length *L* for a 99% chance of detection, while a match length *L* = *v* − 1 gives only 1% detection probability. Note, that compared to ***Equation 2***, we here subtract 1*/*2 in the denominator and *a* = 2 log(99). Then, our model with the exact same parameters (omitting pathogen mutations) results in 18 alleles and *n*_e_ = 16.7, clearly exceeding the number of alleles found by Borghans et al. (2004).

Based on Kimura and Crow (1964), for the above *N* and *µ* the effective number of alleles that can be maintained by heterozygote advantage cannot exceed *n*_e_ = 17.6 at mutation-drift-selection balance. This suggests that the allelic diversity found by Borghans et al. (2004) is likely not limited by the parameters affecting mutation and drift, *µ* and *N*. In contrast, our final number of alleles (being 95% of the maximum), is likely limited by these parameters. To demonstrate that this is indeed the case, we simulate our model with *N* = 10^5^ and *µ* = 5 × 10^−6^, shown in ***Figure S4***. We find well over 100 alleles (*n* = 157 and *n*_e_ = 140). This demonstrates that the ecological parameter values used by Borghans et al. (2004), *m* = 50, *n*_pep_ = 20 and *v* = 7, under our model allows for more than a 20-fold higher allelic diversity.

**Fig. S4.**
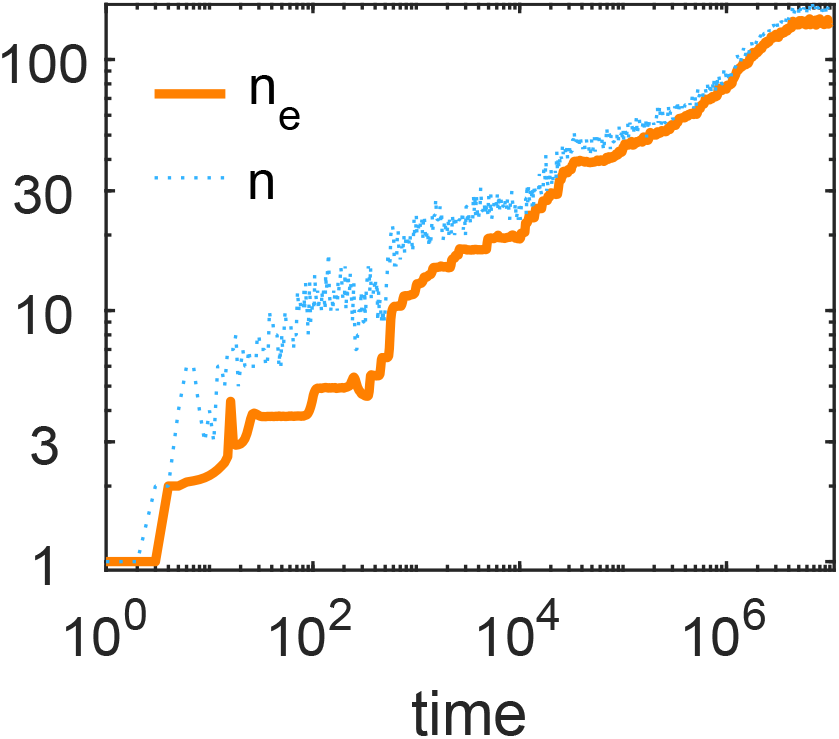
The number of alleles *n* and the effective number of alleles *n*_e_ as a function of time (on a log-log plot) for a simulation run of our bit-string model. Parameters values: *N* = 10^5^ and *µ* = 5 *×* 10^−6^. Other parameters as in Borghans et al. (2004): *v* = 7, *m* = 50, *n*_pep_ = 20.

### 6. Mathematical Analysis of the Gaussian Model: Preliminaries

#### 6.1. Adaptive Dynamics and Invasion Fitness

For the Gaussian model presented in the main part, we investigate with an evolutionary invasion analysis using the adaptive dynamics formalism ***(Metz et al., 1992; Dieckmann and Law, 1996; Geritz et al., 1998)*** whether selection favours a single generalist allele or a polymorphic population. In the language of adaptive dynamics, we ask whether a monomorphic population evolves toward an evolutionary branching point, where two coexisting allelic lineages emerge.

Let us consider a large population of *N* individuals with two segregating alleles ***x***_1_ and ***x***_2_ under Wright-Fisher population dynamics ***(Fisher, 1930; Wright, 1931)***. The allelic frequencies at time *t* are denoted 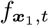 and 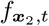.respectively. The recurrence equation for the change of frequency of an allele ***x***_*a*_ ∈ {***x***_1_, ***x***_2_} is then given by

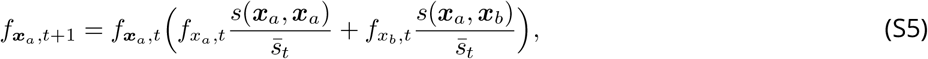

where *s*(***x***_*a*_, ***x***_*b*_) is the survival of an individual carrying the alleles ***x***_*a*_ and ***x***_*b*_ (see ***Equation S12***) and

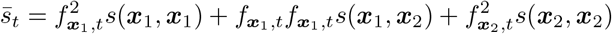

is the population mean survival at time *t*. Note, that the expression within brackets on the right-hand side of ***Equation S5*** describes the marginal fitness of allele ***x***_*a*_.

Consider a resident population carrying allele ***x*** to which a mutant allele ***y*** = ***x*** + ***ϵ*** is introduced. In the limit of a mutant allele-frequency close to zero, its marginal fitness is given by

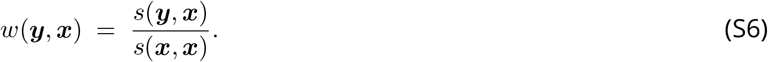

We refer to *w*(***y, x***) as invasion fitness, which is the expected long-term exponential growth rate of an infinitesimally rare mutant allele ***y*** in a resident population with allele ***x (Metz et al., 1992; Metz, 2008)***. Allele ***y*** has a positive probability to invade and increase in frequency if *w*(***y, x***) *>* 1 and disappears otherwise.

We denote the gradient of invasion fitness with respect to the mutant allele ***y***, evaluated at ***y*** = ***x***, with ∇*w*(***x, x***). It has the entries

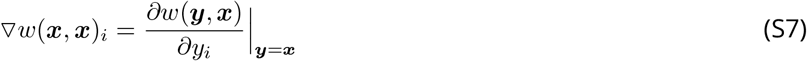

and gives the direction in the *h*-dimensional allelic trait space in which deviations from ***x*** result in the fastest increase of invasion fitness.

If mutations rarely occur, a mutant allele ***y*** will either go extinct or reach an equilibrium frequency before the next mutant appears. If, additionally, ∇*w*(***x, x***) ≠ **0** and mutational effects are sufficiently small (i.e., ***y*** = ***x*** + ***ϵ*** for ***ϵ*** small), then invasion of ***y*** implies extinction of ***x (Dercole and Rinaldi, 2008; Priklopil and Lehmann, 2020)***.

In the limit of small mutational steps, the evolutionary dynamics of an allelic lineage becomes gradual and is given by

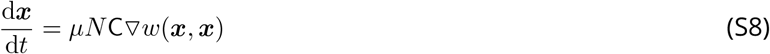

***(Dieckmann and Law, 1996; Champagnat et al., 2006; Durinx et al., 2008; Metz and de Kovel, 2013)***. Here, *µ* is the per-capita mutation probability and C the covariance matrix for the distribution of mutational effects on the trait ***x***.

We note that ***Equation S8*** is structurally similar to the gradient equation of quantitative genetics, which is based on the assumption of weak selection or, equivalently, small genetic variances ***(Lande, 1979; Iwasa et al., 1991; Abrams et al., 1993; Débarre et al., 2014)***. In this case, ***x*** characterizes the mean of the phenotype distribution, the covariance matrix describes the distribution of the standing genetic variation, and the factor *µN* is replaced with a constant.

Allelic trait values ***x*** where ∇*w*(***x, x***) = 0 are of special interest, and such ***x*** are referred to as evolutionarily singular points ***x***^*^. Evolutionarily singular points can be either attractors or repellers of the evolutionary dynamics described by ***Equation S8***. Furthermore, an evolutionarily singular point can be either invadable or uninvadable by nearby mutants. For a resident allele with a one-dimensional trait ***x*** = *x*, a classification of singular strategies is straight forward ***(Geritz et al., 1998)***. Evolutionarily singular points that are not approached, irrespective of whether they are invadable or uninvadable, act as repellers, and we do not expect to ever find resident alleles with such values. Evolutionarily singular strategies that are attractors and uninvadable are endpoints of the evolutionary dynamics. Finally, evolutionarily singular points that are attractors and invadable are known as evolutionary branching points. In this case, any nearby mutant can invade the singular point and coexist with it in a protected dimorphism. Further evolution leads to divergence of the alleles present in the dimorphism. Thus, evolutionary branching points are points in trait space at which diversity emerges ***(Geritz et al., 1998; Rueffler et al., 2006)***.

The classification of singular points becomes more complicated in multivariate trait spaces or when several strategies coexist in an evolutionarily singular point ***(Leimar, 2009; Doebeli, 2011; Geritz et al., 2016)***. First, in multivariate trait spaces or polymorphic populations, whether a singular point is an attractor does not only depend on the direction of the fitness gradient in the vicinity of the singular point but also on the mutational input ***(Leimar, 2009)***. Second, in multivariate trait spaces or polymorphic populations, for evolutionary branching it is necessary that a singular point is an attractor and invadable. However, in the multidimensional case, this is generally not sufficient any more ***(Geritz et al., 2016)***.

In ***Section 7.1***, we show for our model that a unique singular point ***x***^*^ exists. This allele is uninvadable if it is a minimum of *w*(***y, x***^*^) as a function of ***y***. This is the case if the *h*-dimensional Hessian matrix H with entries

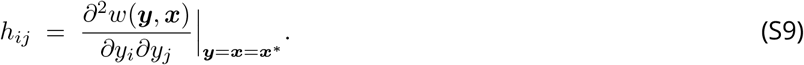

is negative definite ***(Leimar, 2009; Doebeli, 2011)***. In ***Section 7.3*** we derive an explicit expression for H for the fully general case of our model that allows to determine invadability of ***x***^*^ as a function of the positions of the pathogen vectors, the half-saturation constant *K*, and the covariance matrices Σ_*k*_ that determine the shape of the efficiency functions *e*_*k*_.

Whether the singular point ***x***^*^ is an attractor of the evolutionary dynamics can be evaluated based on the Jacobian matrix J of the fitness gradient o*w*(***x***^*^, ***x***^*^) ***(Leimar, 2009)***, which is given by

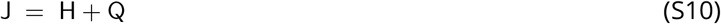

and where Q is the *h*-dimensional matrix of mixed derivatives with entries

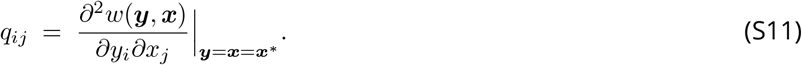

***Leimar (2009)*** shows that if the symmetric part of J, i.e., (J + J^T^)*/*2, is negative definite, then the singular point is an attractor of the evolutionary dynamics described by ***Equation S8*** independent of the mutational covariance matrix C and he refers to this case as strong convergence stability. For the case that the Jacobian matrix is a symmetric negative definite matrix, a stronger result holds, to which he refers to as absolute convergence stability ***(Leimar, 2001, 2009)***. In this case, all conceivable gradualistic, adaptive paths starting near the point ***x***^*^ converge to it. Furthermore, he shows that the condition for absolute convergence stability is equivalent to the existence of a function *g*(***x***) having a maximum at ***x***^*^ and a positive function *α*(***x***) such that the gradient of invasion fitness can be expressed as

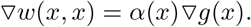

In ***Section 7.2***, we show for our model that ***x***^*^ is indeed absolutely convergence stable.

For the case of two-dimensional trait spaces, results in ***Geritz et al. (2016)*** allow us to conclude that if ***x***^*^ is invadable, then it is indeed an evolutionary branching point. For trait spaces of dimension three or higher, whether convergence stability and invadability imply evolutionary branching is an open problem ***(Geritz et al., 2016)***. Individual-based simulations indicate, however, that for our model this is indeed the case.

#### 6.2. Model Description

In this section, we describe the model ingredients. Survival *s*(***x***_*a*_, ***x***_*b*_) of a genotype carrying alleles ***x***_*a*_ and ***x***_*b*_ is a saturating function of condition *c* and described by the well known Michaelis-Menten equation

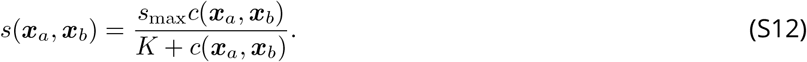

Here, the half saturation constant *K* gives the condition *c* at which half of the maximum survival is reached and *s*_max_ is the maximum survival probability that is approached when *c* becomes large.

The condition of a genotype is given by

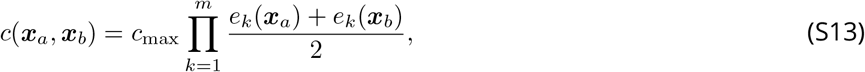

where *c*_max_ is the condition of a hypothetical individual with perfect defence against all *m* pathogens and *e*_*k*_(***x***) is the efficiency of an allele’s MHC molecule against pathogen *k* in an environment with *m* pathogens.

Without loss of generality, *c*(***x***^*^, ***x***^*^) is standardized to 1 (by choosing *c*_max_ in ***Equation S13*** appropriately). This is helpful because it allows us to choose an interval of *K*-values where individuals homozygous for the generalist allele have either a condition in the range where survival changes rapidly (*K >>* 1) or slowly (*K <<* 1) with condition. In ***Figure S3*** and ***4***, the x-axis can be translated into survival *s*(***x***^*^, ***x***^*^) of the generalist genotype using ***Equation S12***, which then varies between 0.01 for *K* = 10^−2^ and 0.99 for *K* = 10^2^.

We assume that the efficiencies *e*_*k*_(***x***) of inducing immune defence against the *m* different pathogens are traded off. This trade-off emerges by describing the efficiencies against different pathogens with multivariate Gaussian functions (see ***Figure 1***) that have pathogen-dependent optima,

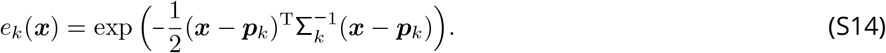

These function describes how the efficiency of an allele characterized by the *h*-dimensional vector ***x*** decreases with increasing distance from the pathogen vector ***p***_*k*_. The closer an allelic trait vector is to a pathogen vector, the higher is the efficiency of the MHC molecule against that pathogen. The magnitude of the decrease in efficiency with increasing distance to the *k*th pathogen is determined by the shape and width of the Gaussian function as determined by the *h*-dimensional covariance matrix Σ_*k*_.

In the main part, we consider the special case of rotationally symmetric Gaussian functions *e*_*k*_(***x***). These matrices are thus specified by an inverse matrix-covariance matrix 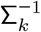 (see ***Equation S14***) that takes the form of a scalar matrix, that is, a scalar multiple of the identity matrix *I*. Furthermore, we assume that all Gaussians are of equal width. Hence, we have a common scalar for all Gaussians that we denote with *v*^2^, that is, *v* is the inverse of the width of the Gaussian function. We refer to *v* as virulence (see ***Section 7.5***, below).

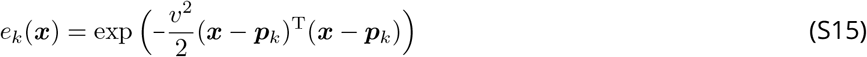

### 7. Analytical Results for the Gaussian Model

#### 7.1. The Evolutionarily Singular Point

In this section, we analyse the evolutionary dynamics of a monomorphic resident population in full generality. By subsequently applying several symmetry assumptions, we then derive the analytical results presented in the main text (see ***Section 7.4*** and ***7.5***). Invasion fitness of a rare mutant allele ***y*** in a resident population with allele ***x*** is given by its marginal fitness,

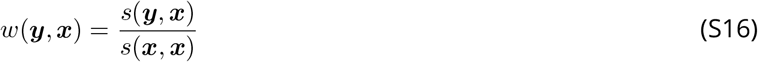

(see derivation of ***Equation S6***). Note, that *s*_max_ cancels out. It is therefore omitted from all further calculations. The direction of the evolutionary dynamics is governed by the selection gradient. Its *i*th entry calculates to

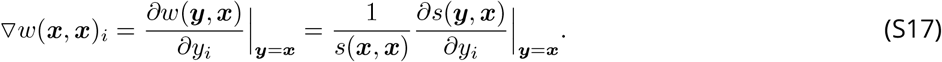

Using the definitions for *s* (***Equation S12***) and *c* (***Equation S13***) and their derivatives,

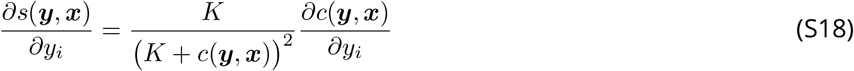

and

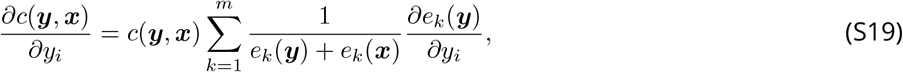

where ***Equation S19*** is obtained by applying the generalised product rule

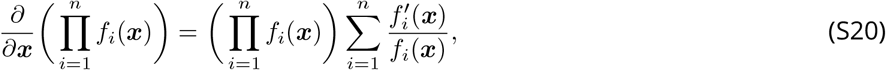

we obtain

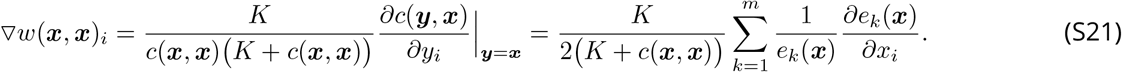

In the next step, we calculate the derivative of the function *e*_*k*_(***x***) (***Equation S14***). Applying the chain rule and simplifying results in

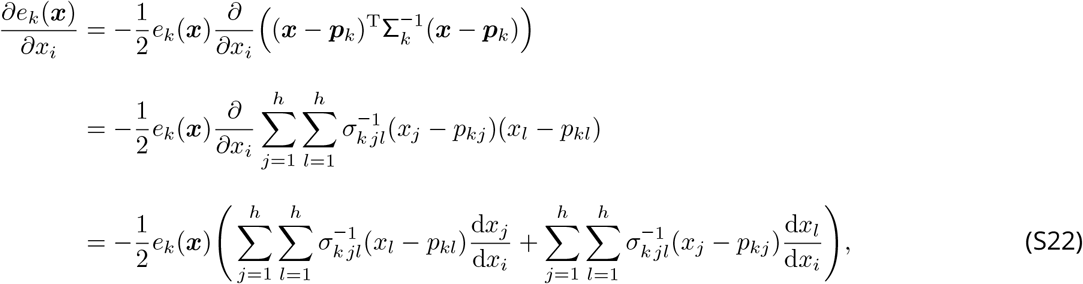

where the entries of the matrix 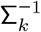 are denoted by 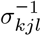. Using that 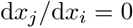 for *i ≠ j* and 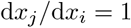 for *i* = *j* this further simplifies to

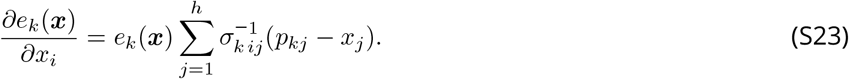

Substituting ***Equation S23*** into ***Equation S21*** finally results in

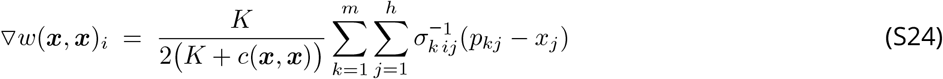

and

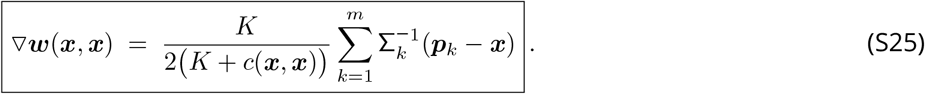

As mentioned in ***Section 6.1***, singular points ***x***^*^ are allelic trait vectors for which ***Equation S25*** equals zero. From ***Equation S25*** follows that in our model singular points have to fulfil

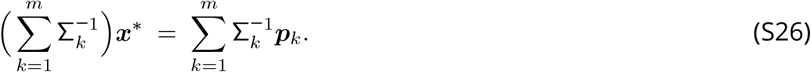

Solving for ***x***^*^ yields

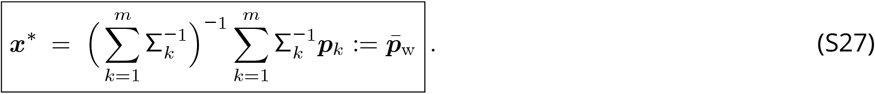

Thus, the unique singular point ***x***^*^ equals the arithmetic mean of the pathogen vectors ***p***_1_, … ***p***_*m*_, each weighted by the inverse of their Gaussian covariance matrices Σ_1_, …, Σ_*m*_. For a one dimensional trait space (*h* = 1) this simplifies to

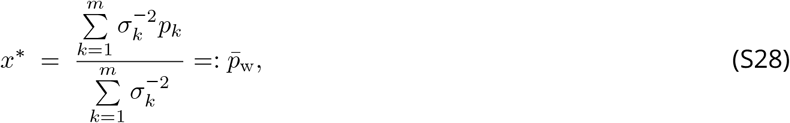

which is the well-known weighted average for scalars. If Σ_1_ = … = Σ_*m*_, then ***Equation S27*** simplifies to the arithmetic mean pathogen vector

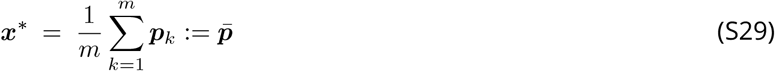

as stated in the main text.

#### 7.2. Absolute Convergence Stability

Below, we prove that the unique singular point ***x***^*^ (***Equation S27***) is absolutely convergence stable. To this end, we first demonstrate that the gradient of invasion fitness can be expressed as ∇*w*(*x, x*) = *α*(*x*) ∇*c*(*x, x*), where *α*(*x*) is a positive function, and then show that ***x***^*^ is a maximum of *c*(*x, x*).

##### 7.2.1. Gradient of Condition

An individual homozygote for ***x*** has a condition given by

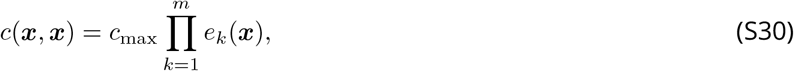

and the *i*th entry of the gradient ∇*c*(*x, x*) is given by

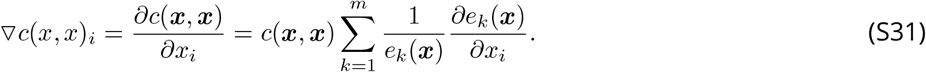

Substituting ***Equation S23*** into ***Equation S31*** gives

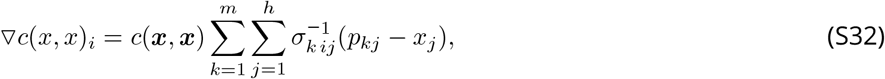

and

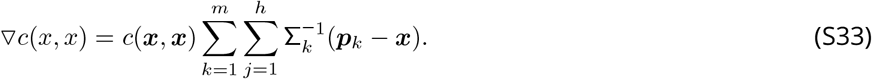

By comparing ***Equation S33*** with ***Equation S25*** we see that

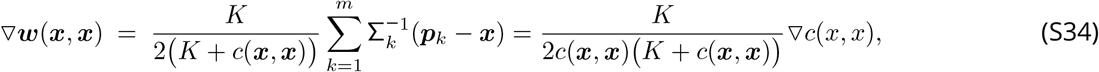

where the fraction before ∇*c*(*x, x*) is positive. Thus, ∇*w*(*x, x*) = *α*(*x*) ∇*c*(*x, x*).

##### 7.2.2. Hessian Matrix of Condition

The Hessian matrix of a homozygote individual’s condition, evaluated at the singular point, is given by the second order partial derivative of *c*(***x***^*^, ***x***^*^) with respect to the *i*th and *j*th entry of ***x***^*^. We obtain this by differentiating ***Equation S33*** with respect of *x*_*j*_, evaluated at ***x*** = ***x***^*^, resulting in

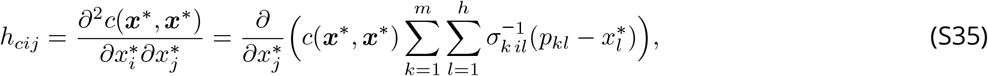

and applying the product rule and using that the first derivative at ***x***^*^ is zero gives

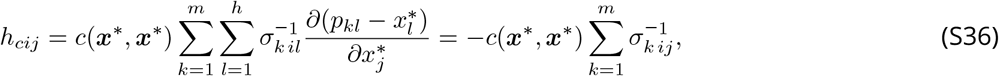

where the last simplification uses that d*x*_*j*_*/*d*x*_*i*_ = 0 for *i* ≠ *j* and d*x*_*j*_*/*d*x*_*i*_ = 1 for *i* = *j*, and

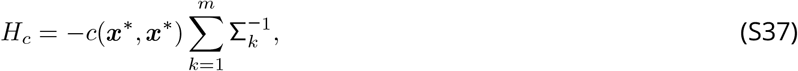

which is always negative definite. Thus, *c*(***x***^*^, ***x***^*^) is a maximum and ***x***^*^ is absolute convergence stable.

#### 7.3. Derivation of the Hessian Matrix of Invasion Fitness

As stated in ***Equation S9***, the entries of the Hessian matrix of invasion fitness are given by

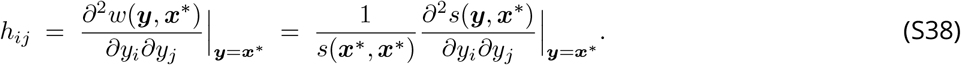

The second derivative of the function *s* is obtained by differentiating ***Equation S18*** with respect to *y*_*j*_, resulting in

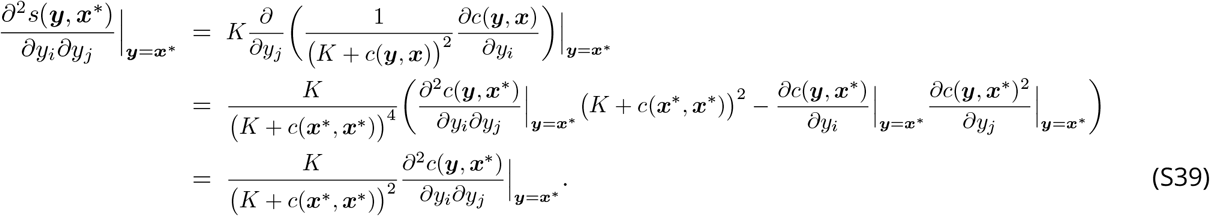

In the final simplification step we use the conclusion drawn from ***Equation S21*** that 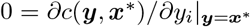 and therefore the term after the minus sign disappears.

The second derivative for the function *c* is obtained by differentiating ***Equation S19*** with respect to *y*_*j*_, resulting in

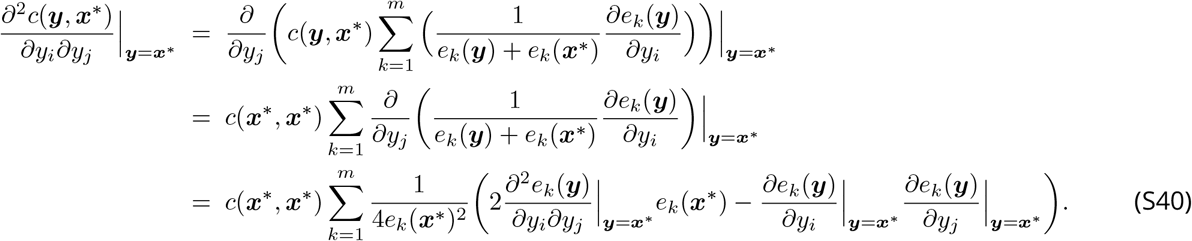

Here, the one but last simplification step again follows from the fact that 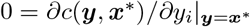.

The second derivative for the function *e*_*k*_ is obtained by differentiating ***Equation S23*** with respect to *y*_*j*_, resulting in

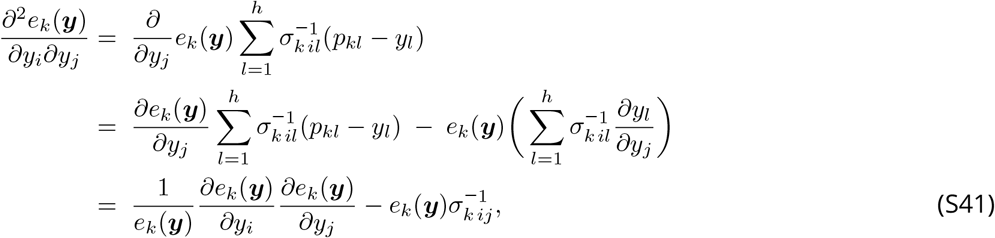

where the last simplification uses that d*y*_*j*_*/*d*y*_*i*_ = 0 for *i* ≠ *j* and d*y*_*j*_*/*d*y*_*i*_ = 1 for *i* = *j*.

By recursively substituting ***Equations S39-S41*** into ***Equation S9*** we obtain

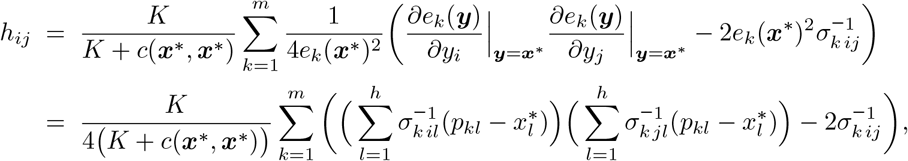

where in the last step we substituted ***Equation S23***. This result can be rewritten as a matrix

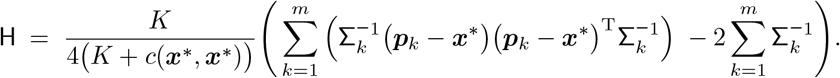

Finally, substituting ***x***^*^ with ***Equation S27***, we obtain

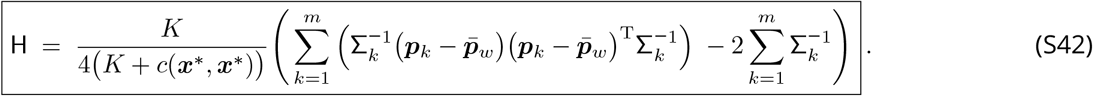

#### 7.4. Special Case: Identically Shaped Gaussian Efficiency Functions

For the special case that the Gaussian covariance matrices Σ_*k*_ are equal (Σ_1_ = Σ_2_ = … = Σ_*m*_ = Σ_G_), fulfilled in ***Figure S3B***, the Hessian matrix simplifies to

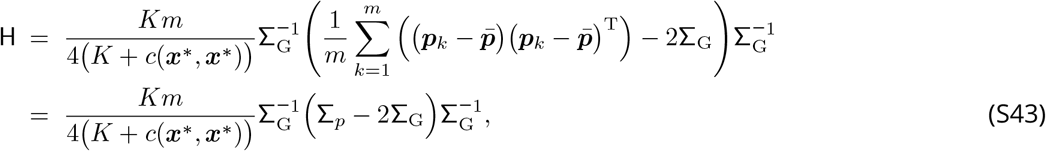

where we used that 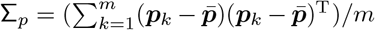 is the covariance matrix of the positions ***p***_*k*_ of the pathogen vectors. Since the fraction in ***Equation S43*** is positive and Σ_G_^−1^ is positive definite, it follows that the definiteness of H is given by the definiteness of the matrix Σ_*p*_ − 2Σ_G_. Hence, the singular point ***x***^*^ is uninvadable whenever

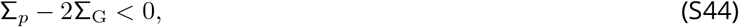

where the inequality indicates negative definiteness.

#### 7.5. Special Case: Maximal Symmetry

For the even more special case that 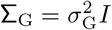 and that the pathogen vectors are arranged symmetrically, resulting in 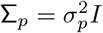,we obtain

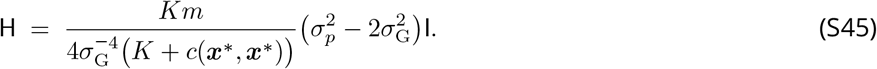

In this case, the singular point ***x***^*^ is a branching point whenever 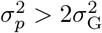. Assuming that all *m* pathogen vectors have an equal distance *d* to each other implies that they are arranged in an *m* − 1-dimensional regular simplex (see ***Figure 1***). For this case, 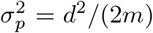,and we obtain the branching condition 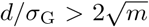.Assuming pathogen distance *d* = 1 and substitute 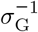 as virulence *v*, we obtain

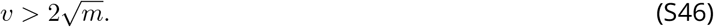

## Notes

### Competing Interest Statement

The authors have declared no competing interest.

### Summary of Updates

Manuscript revised for enhanced clarity with an expanded discussion on pathogen co-evolution and MHC diversity.

## References

Abbas, A. K., A. H. Lichtman, and S. Pillai. 2014. Basic immunology: functions and disorders of the immune system. Elsevier Health Sciences.

Alizon, S., A. Hurford, N. Mideo, and M. Van Baalen. 2009. Virulence evolution and the trade-off hypothesis: history, current state of affairs and the future. Journal of evolutionary biology 22:245–259.

Allison, A. C. 1954. Protection afforded by sickle-cell trait against subtertian malarial infection. British medical journal 1:290.

Anderson, R. M., and R. M. May. 1982. Coevolution of hosts and parasites. Parasitology 85:411–426.

Apanius, V., D. Penn, P. R. Slev, L. R. Ruff, and W. K. Potts. 1997. The nature of selection on the major histocompatibility complex. Critical Reviews in Immunology 17.

Borghans, J. A., J. B. Beltman, and R. J. De Boer. 2004. MHC polymorphism under host-pathogen coevolution. Immunogenetics 55:732–739.

Castric, V., and X. Vekemans. 2004. Plant self-incompatibility in natural populations: a critical assessment of recent theoretical and empirical advances. Molecular Ecology 13:2873–2889.

Chappell, P. E., E. K. Meziane, M. Harrison, Ł. Magiera, C. Hermann, L. Mears, A. G. Wrobel, C. Durant, L. L. Nielsen, S. Buus, et al. 2015. Expression levels of MHC class I molecules are inversely correlated with promiscuity of peptide binding. Elife 4.

Cortazar-Chinarro, M., S. Meurling, L. Schroyens, M. Siljestam, A. Richter-Boix, A. Laurila, and J. Höglund. 2022. Major histocompatibility complex variation and haplotype associated survival in response to experimental infection of two bd-gpl strains along a latitudinal gradient. Frontiers in Ecology and Evolution page 653.

de Boer, R. J., J. A. Borghans, M. Van Boven, C. Keşmir, and F. J. Weissing. 2004. Heterozygote advantage fails to explain the high degree of polymorphism of the MHC. Immunogenetics 55:725–731.

Ding, G., M. Hasselmann, J. Huang, J. Roberts, B. P. Oldroyd, and R. Gloag. 2021. Global allele polymorphism indicates a high rate of allele genesis at a locus under balancing selection. Heredity 126:163–177.

Doebeli, M. 2011. Adaptive Diversification (MPB-48). Princeton University Press.

Doherty, P. C., and R. M. Zinkernagel. 1975. Enhanced immunological surveillance in mice heterozygous at the h-2 gene complex. Nature 256:50–52.

Duncan, W. R., E. K. Wakeland, and J. Klein. 1979. Histocompatibility-2 system in wild mice. Immunogenetics 9:261–272.

Eizaguirre, C., and T. Lenz. 2010. Major histocompatibility complex polymorphism: dynamics and consequences of parasite-mediated local adaptation in fishes. Journal of Fish Biology 77:2023–2047.

Eizaguirre, C., T. L. Lenz, M. Kalbe, and M. Milinski. 2012. Divergent selection on locally adapted major histocompatibility complex immune genes experimentally proven in the field. Ecology letters 15:723–731.

Ejsmond, M. J., and J. Radwan. 2015. Red queen processes drive positive selection on major histocompatibility complex (MHC) genes. PLoS Comput Biol 11.

Ejsmond, M. J., J. Radwan, A. Ejsmond, T. Gaczorek, and W. Babik. 2023. Adaptive immune response selects for postponed maturation and increased body size. Functional Ecology 37:2883–2894.

Fisher, R. A. 1930. The genetical theory of natural selection. Oxford University Press.

Frank, S., and P. Schmid-Hempel. 2008. Mechanisms of pathogenesis and the evolution of parasite virulence. Journal of evolutionary biology 21:396–404.

Frank, S. A. 1996. Models of parasite virulence. The Quarterly review of biology 71:37–78.

Froeschke, G., and S. Sommer. 2005. MHC class II DRB variability and parasite load in the striped mouse (Rhabdomys pumilio) in the southern Kalahari. Molecular biology and evolution 22:1254–1259.

Froeschke, G., and S. Sommer. 2012. Insights into the complex associations between MHC class II DRB polymorphism and multiple gastrointestinal parasite infestations in the striped mouse. PLoS One 7.

Geritz, S. A. H., É. Kisdi, G. Meszéna, and J. A. J. Metz. 1998. Evolutionarily singular strategies and the adaptive growth and branching of the evolutionary tree. Evolutionary Ecology 12:35–57.

Gillespie, J. H. 2004. Population Genetics: A Concise Guide. The Johns Hopkins University Press.

Gould, S. J., J. E. Hildreth, and A. M. Booth. 2004. The evolution of alloimmunity and the genesis of adaptive immunity. The Quarterly review of biology 79:359–382.

Hedrick, P. W. 1999. Balancing selection and MHC. Genetica 104:207–214.

Hedrick, P. W. 2002. Pathogen resistance and genetic variation at MHC loci. Evolution 56:1902–1908.

Hedrick, P. W. 2012. What is the evidence for heterozygote advantage selection? Trends in ecology & evolution 27:698–704.

Hughes, A. L., and M. Nei. 1989. Nucleotide substitution at major histocompatibility complex class II loci: evidence for over-dominant selection. Proceedings of the National Academy of Sciences 86:958–962.

Jeffery, K. J., and C. R. Bangham. 2000. Do infectious diseases drive MHC diversity? Microbes and Infection 2:1335–1341.

Kekäläinen, J., J. A. Vallunen, C. R. Primmer, J. Rättyä, and J. Taskinen. 2009. Signals of major histocompatibility complex over-dominance in a wild salmonid population. Proceedings of the Royal Society of London B: Biological Sciences.

Kimura, M., and J. F. Crow. 1964. The number of alleles that can be maintained in a finite population. Genetics 49:725–738.

Kisdi, É., and S. A. H. Geritz. 1999. Adaptive dynamics in allele space: evolution of genetic polymorphism by small mutations in a heterogeneous environment. Evolution pages 993–1008.

Klitz, W., P. Hedrick, and E. J. Louis. 2012. New reservoirs of hla alleles: pools of rare variants enhance immune defense. Trends in Genetics 28:480–486.

Lau, Q., Y. Yasukochi, and Y. Satta. 2015. A limit to the divergent allele advantage model supported by variable pathogen recognition across hla-drb1 allele lineages. Tissue Antigens 86:343–352.

Lenz, T. L. 2011. Computational prediction of MHC II-antigen binding supports divergent allele advantage and explains trans-species polymorphism. Evolution 65:2380–2390.

Levins, R. 1968. Evolution in changing environments: some theoretical explorations. 2. Princeton University Press.

Lewontin, R., L. Ginzburg, and S. Tuljapurkar. 1978. Heterosis as an explanation for large amounts of genic polymorphism. Genetics 88:149–169.

Lighten, J., A. S. Papadopulos, R. S. Mohammed, B. J. Ward, I. G. Paterson, L. Baillie, I. R. Bradbury, A. P. Hendry, P. Bentzen, and C. Van Oosterhout. 2017. Evolutionary genetics of immunological supertypes reveals two faces of the red queen. Nature communications 8:1294.

Linnenbrink, M., M. Teschke, I. Montero, M. Vallier, and D. Tautz. 2018. Meta-populational demes constitute a reservoir for large MHC allele diversity in wild house mice (Mus musculus). Frontiers in zoology 15:15.

Loiseau, C., R. Zoorob, A. Robert, O. Chastel, R. Julliard, and G. Sorci. 2011. Plasmodium relictum infection and MHC diversity in the house sparrow (Passer domesticus). Proceedings of the Royal Society of London B: Biological Sciences 278:1264–1272.

Maruyama, T., and M. Nei. 1981. Genetic variability maintained by mutation and overdominant selection in finite populations. Genetics 98:441–459.

McClelland, E. E., D. J. Penn, and W. K. Potts. 2003. Major histocompatibility complex heterozygote superiority during coinfection. Infection and immunity 71:2079–2086.

Metz, J. A. J., R. M. Nisbet, and S. A. H. Geritz. 1992. How should we define ‘fitness’ for general ecological scenarios? Trends in Ecology & Evolution 7:198–202.

Oliver, M., S. Telfer, and S. Piertney. 2009. Major histocompatibility complex (MHC) heterozygote superiority to natural multi-parasite infections in the water vole (arvicola terrestris). Proceedings of the Royal Society of London B: Biological Sciences 276:1119–1128.

Penn, D. J. 2002. The scent of genetic compatibility: sexual selection and the major histocompatibility complex. Ethology 108:1–21.

Penn, D. J., K. Damjanovich, and W. K. Potts. 2002. MHC heterozygosity confers a selective advantage against multiple-strain infections. Proceedings of the National Academy of Sciences 99:11260–11264.

Pierini, F., and T. L. Lenz. 2018. Divergent allele advantage at human MHC genes: signatures of past and ongoing selection. Molecular Biology and Evolution 35:2145–2158.

Proulx, S. R., and P. C. Phillips. 2006. Allelic divergence precedes and promotes gene duplication. Evolution 60:881–892.

Robinson, J., L. A. Guethlein, N. Cereb, S. Y. Yang, P. J. Norman, S. G. Marsh, and P. Parham. 2017. Distinguishing functional polymorphism from random variation in the sequences of> 10,000 hla-a,-b and-c alleles. PLoS Genetics 13:e1006862.

Schmid-Hempel, P. 2021. Evolutionary parasitology: the integrated study of infections, immunology, ecology, and genetics. Oxford University Press.

Sellis, D., D. J. Kvitek, B. Dunn, G. Sherlock, and D. A. Petrov. 2016. Heterozygote advantage is a common outcome of adaptation in saccharomyces cerevisiae. Genetics 203:1401–1413.

Sheftel, H., P. Szekely, A. Mayo, G. Sella, and U. Alon. 2018. Evolutionary trade-offs and the structure of polymorphisms. Philosophical Transactions of the Royal Society B: Biological Sciences 373:20170105.

Sommer, S. 2005. The importance of immune gene variability (MHC) in evolutionary ecology and conservation. Frontiers in Zoology 2:16.

Spencer, H. G., and R. W. Marks. 1988. The maintenance of single-locus polymorphism. I. Numerical studies of a viability selection model. Genetics 120:605–613.

Spurgin, L. G., and D. S. Richardson. 2010. How pathogens drive genetic diversity: MHC, mechanisms and misunderstandings. Proceedings of the Royal Society of London B: Biological Sciences 277:979–988.

Stefan, T., L. Matthews, J. M. Prada, C. Mair, R. Reeve, and M. J. Stear. 2019. Divergent allele advantage provides a quantitative model for maintaining alleles with a wide range of intrinsic merits. Genetics 212:553–564.

Stoffels, R. J., and H. G. Spencer. 2008. An asymmetric model of heterozygote advantage at major histocompatibility complex genes: degenerate pathogen recognition and intersection advantage. Genetics 178:1473–1489.

Takahata, N., and M. Nei. 1990. Allelic genealogy under overdominant and frequency-dependent selection and polymorphism of major histocompatibility complex loci. Genetics 124:967–978.

Thompson, W. W., D. K. Shay, E. Weintraub, L. Brammer, C. B. Bridges, N. J. Cox, and K. Fukuda. 2004. Influenza-associated hospitalizations in the united states. Jama 292:1333–1340.

Trotter, M. V., and H. G. Spencer. 2008. The generation and maintenance of genetic variation by frequency-dependent selection: constructing polymorphisms under the pairwise interaction model. Genetics 180:1547–1557.

Trotter, M. V., and H. G. Spencer. 2013. Models of frequency-dependent selection with mutation from parental alleles. Genetics 195:231–242.

Wakeland, E. K., S. Boehme, J. X. She, C.-C. Lu, R. A. McIndoe, I. Cheng, Y. Ye, and W. K. Potts. 1990. Ancestral polymorphisms of MHC class II genes: divergent allele advantage. Immunologic Research 9:115–122.

Wegner, K. 2008. Historical and contemporary selection of teleost MHC genes: did we leave the past behind? Journal of Fish Biology 73:2110–2132.

Wilder, S. M., D. Raubenheimer, and S. J. Simpson. 2016. Moving beyond body condition indices as an estimate of fitness in ecological and evolutionary studies. Functional Ecology 30:108–115.

Wright, S. 1931. Evolution in Mendelian populations. Genetics 16:97–159.

Wright, S. 1939. The distribution of self-sterility alleles in populations. Genetics 24:538.

Wright, S. 1966. Polyallelic random drift in relation to evolution. Proceedings of the National Academy of Sciences 55:1074–1081.

Zhou, F., T. Yu, R. Du, G. Fan, Y. Liu, Z. Liu, J. Xiang, Y. Wang, B. Song, X. Gu, et al. 2020. Clinical course and risk factors for mortality of adult inpatients with covid-19 in wuhan, china: a retrospective cohort study. The lancet 395:1054–1062.

## References

Abrams, P. A., H. Matsuda, and Y. Harada. 1993. Evolutionarily unstable fitness maxima and stable fitness minima of continuous traits. Evolutionary Ecology 7:465–487.

Champagnat, N., R. Ferrière, and S. Méléard. 2006. Unifying evolutionary dynamics: from individual stochastic processes to macroscopic models. Theoretical Population Biology 69:297–321.

Débarre, F., S. L. Nuismer, and M. Doebeli. 2014. Multidimensional (co) evolutionary stability. The American Naturalist 184:158–171.

Dercole, F., and S. Rinaldi. 2008. Analysis of evolutionary processes: the adaptive dynamics approach and its applications: the adaptive dynamics approach and its applications. Princeton University Press.

Dieckmann, U., and R. Law. 1996. The dynamical theory of coevolution: a derivation from stochastic ecological processes. Journal of Mathematical Biology 34:579–612.

Doebeli, M. 2011. Adaptive Diversification (MPB-48). Princeton University Press.

Durinx, M., J. A. J. H. Metz, and G. Meszéna. 2008. Adaptive dynamics for physiologically structured population models. Journal of Mathematical Biology 56:673–742.

Fisher, R. A. 1930. The genetical theory of natural selection. Oxford University Press.

Geritz, S. A. H., J. A. J. Metz, and C. Rueffler. 2016. Mutual invadability near evolutionarily singular strategies for multivariate traits, with special reference to the strongly convergence stable case. Journal of Mathematical Biology 72:1081–1099.

Iwasa, Y., A. Pomiankowski, and S. Nee. 1991. The evolution of costly mate preferences ii. the “handicap” principle. Evolution 45:1431–1442.

Johnson, N. L., S. Kotz, and N. Balakrishnan. 1994. Continuous univariate distributions, volume 1. John wiley & sons.

Lande, R. 1979. Quantitative genetic analysis of multivariate evolution, applied to brain: body size allometry. Evolution 33:402–416.

Leimar, O. 2001. Evolutionary change and darwinian demons. Selection 2:65–72.

Leimar, O. 2009. Multidimensional convergence stability. Evolutionary Ecology Research 11:191–208.

Metz, J. A. J. 2008. Fitness. Pages 1599–1612 in S. E. Jorgensen and B. Fath, eds. Encyclopedia of ecology. Oxford: Elsevier, available online as IIASA Interim Report IR-06-061.

Metz, J. A. J., and C. G. F. de Kovel. 2013. The canonical equation of adaptive dynamics for Mendelian diploids and haplo-diploids. Interface Focus 3.

Priklopil, T., and L. Lehmann. 2020. Invasion implies substitution in ecological communities with class-structured populations. Theoretical Population Biology 134:36–52.

Rueffler, C., T. J. M. Van Dooren, O. Leimar, and P. A. Abrams. 2006. Disruptive selection and then what? Trends in Ecology & Evolution 21:238–245.

Wright, S. 1931. Evolution in Mendelian populations. Genetics 16:97–159.

